# Deleterious mutations and selection for sex in spatially structured, diploid populations

**DOI:** 10.1101/2025.01.22.634382

**Authors:** Louise Fouqueau, Denis Roze

## Abstract

By increasing local genetic drift and generating inbreeding, population spatial structure may have important effects on the evolutionary benefits of sexual reproduction. In this article, we consider a population structured according to the island model, and use two and three-locus analytical models and multilocus simulations to explore the selective forces acting on a modifier locus affecting the rate of sexual reproduction of facultatively sexual organisms, in the presence of recurrent deleterious mutations. The results show that population structure and selection combine to generate a local excess of heterozygotes at selected loci (negative *F*_IS_), for both partially recessive and partially dominant deleterious alleles. The linkage disequilibrium between deleterious alleles may be either negative or positive depending on their dominance coefficient, the degree of population structure and the rate of sex. These genetic associations combine with many other ones to generate indirect selection at the sex modifier locus, generally favoring intermediate rates of sex even when sex entails direct fitness costs. Multilocus simulations show that the equilibrium rate of sex increases moderately as the degree of population structure increases. However, population structure may also prevent the irreversible spread of asexual mutants when the cost of sex is strong.

## INTRODUCTION

Several theories have been proposed to explain the widespread occurrence of sexual reproduction among eukaryotic species (Otto, 2009; Hartfield and Keightley, 2012). These theories correspond to different scenarios under which the effect of sex on the distribution of fitness among offspring bears an advantage, by increasing either the mean fitness or the variance in fitness among offspring (Otto and Lenormand, 2002; Agrawal, 2006; Sharp and Otto, 2016). In haploid species, this effect of sex is due to genetic recombination occurring during meiosis, that tends to break linkage disequilibrium (LD) between loci (Felsenstein, 1974). Epistatic interactions among selected loci represent a possible source of LD, and several models have shown that recombination can be favored when the sign of epistasis fluctuates over time (which may occur in models of coevolution between hosts and parasites, e.g., Peters and Lively, 1999; Otto and Nuismer, 2004; Gandon and Otto, 2007; Salathé et al., 2009), or in the presence of synergistic epistasis between deleterious mutations (in which case recombination decreases the mean fitness of offspring, but increases the variance in fitness and the efficiency of selection, e.g., Charlesworth, 1990; Barton, 1995a). Another possible source of LD stems from interference (or Hill-Robertson effect) occurring between selected loci in finite populations (Hill and Robertson, 1966; Barton, 1995b; Otto, 2021), that tends to favor higher recombination rates (Otto and Barton, 1997; Barton and Otto, 2005; Keightley and Otto, 2006; Roze, 2021).

In diploid species, sex may also affect the fitness of offspring through their degree of heterozygosity: in particular, a single round of random mating in an infinite population suffices to bring the population to Hardy-Weinberg equilibrium (HWE), dissipating any excess or deficit of heterozygotes that may exist among the parents. This ‘segregation’ effect of sex may be favored or disfavored when selection (or any other force) tends to generate systematic deviations from HWE. This is the case in the presence of dominance interactions between alleles at selected loci, defined as a deviation from multiplicative effects (on fitness) of the two alleles present in a diploid individual. In particular, when deleterious alleles are recessive on a multiplicative scale 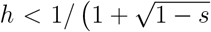 if *AA, Aa* and *aa* individuals have fitnesses 1, 1 − *sh* and 1 − *s*), selection generates an excess of heterozygotes relative to HWE (Chasnov, 2000; Otto, 2003). In this case, sex increases the efficiency of selection by increasing the frequency of homozygotes among offspring, and thus reduces the mutation load. In an infinite, randomly mating population, this reduction in load is typically rather small, however, unless *h* is close to zero (Chasnov, 2000). Furthermore, in a facultatively sexual population in which individuals produce a fraction of their offspring clonally and the remaining fraction through sexual reproduction, genotypes coding for higher rates of sex tend to produce offspring with lower mean fitness (due to the production of unfit homozygotes). Under random mating, this immediate cost of segregation prevents high rates of sex to evolve unless selection is close to multiplicative (*h* slightly below 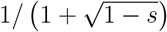), in particular when deleterious alleles have weak fitness effects (Uyenoyama and Bengtsson, 1989; Otto, 2003).

Finite population size can amplify the difference in mutation load between obligately sexual and obligately clonal populations, particularly in regimes where mutations may fix in the heterozygous state in clonal populations, while they are still efficiently selected against in sexual populations due to segregation (Haag and Roze, 2007). Furthermore, finite population size tends to generate an excess of heterozygotes relative to HWE, even in the absence of selection; this effect is small under obligate sex, but becomes stronger as the rate of clonality increases (Roze and Michod, 2010; Hartfield et al., 2016). In the case of a deleterious allele maintained at mutation-selection balance, this effect is stronger than deviations from HWE generated by selection roughly when 1*/* (2*N*_e_) is larger than *u* |1 − 2*h*| */h*, where *N*_e_ is the effective population size and *u* the (per locus) deleterious mutation rate (Roze and Michod, 2010). This can favor sex in the presence of dominant deleterious alleles, as segregation then increases both the mean fitness and the variance in fitness among offspring by producing more homozygous offspring. However, in the more realistic case of partially recessive deleterious mutations, one obtains again that, in the absence of any direct cost of sex, high rates of sex can be favored only mutations are slightly recessive (Roze and Michod, 2010). When a direct cost of sex is introduced, multilocus simulations show that populations often evolve towards obligate clonality when *h* ≤ 0.3 (Figure 7 in Roze and Michod, 2010).

Most populations present some form of geographical structure, due to the limited dispersal abilities of organisms. Spatial population structure has several evolutionary consequences; in particular: (i) it may lead to adaptations to local environmental conditions, (ii) it may increase the effects of drift within local populations, and (iii) it generates inbreeding (at the level of the whole metapopulation) when mating occurs locally. These three effects have in turn important consequences on selection for sex. In spatially heterogeneous environments, local adaptation can increase the parameter range under which recombination and segregation are favored (Pylkov et al., 1998; Lenormand and Otto, 2000; Agrawal, 2009), while it may also favor the spread of asexual lineages at species range limits (Peck et al., 1998; Fouqueau and Roze, 2021). Increased local drift enhances the strength of the Hill-Robertson effect, and thus increases selection for recombination (Martin et al., 2006). It also reduces the fixation probability of asexual mutants, since asexual lineages may accumulate deleterious mutations by Muller’s ratchet before reaching fixation at the metapopulation level (Peck et al., 1999; Salathé et al., 2006; Hartfield et al., 2012). Finally, inbreeding tends to increase the efficiency of selection against deleterious alleles (by increasing homozygosity), thus decreasing the mutation load of sexual populations relative to asexual ones (Agrawal and Chasnov, 2001; Haag and Roze, 2007).

While these results suggest that population structure should generally enhance selection for sex, several questions remain open. In particular, most models on the evolution of sex in structured populations either compare the equilibrium mutation load of sexual and asexual populations (e.g., Agrawal and Chasnov, 2001; Haag and Roze, 2007) or consider the spread of an asexual mutant in an obligately sexual population (Peck et al., 1999; Salathé et al., 2006; Hartfield et al., 2012), and one may wonder how population structure may affect the evolutionarily stable rate of sex in facultatively sexual species. This should depend not only on the long-term effect of sex on the equilibrium mutation load, but also on the immediate effect of sex on offspring fitness, and it is not obvious how population structure will affect this immediate effect and its importance relative to longer term benefits. Furthermore, most previous models either assume haploid organisms (in which sex only affects the fitness of offspring through recombination) or neglect the effect of recombination between selected loci (focusing on the effect of segregation), so that the relative importance of recombination and segregation on selection for sex often stays unclear.

In this article, we explore the selective forces acting on a modifier locus affecting the rate of sex in a facultatively sexual population, structured according to the island model with a very large number of demes. Analytical results are derived in the case of weak population structure (*Nd* ≫ 1, where *N* is deme size and *d* the dispersal rate) and when the number of demes tends to infinity, assuming either a single selected locus undergoing recurrent deleterious mutation (two-locus model, capturing selection for sex through segregation), or two selected loci (three-locus model, also including the effect of recombination). In the two-locus model, indirect selection for sex results from an excess of heterozygosity within demes at the selected locus (resulting from the combined effects of selection and population structure), and from various other effects generated by the variance among demes in LD between the modifier and the selected locus. The three-locus model shows that the LD between selected loci may be either positive or negative depending on the degree of population structure, rate of sex and dominance coefficient of deleterious alleles, while selection for sex also depends on other forms of interactions between selected loci, that generally tend to favor higher rates of sex. The results are then extrapolated to the case of a very large number of selected loci, and compared with individual-based simulation results. Overall, our results show that population structure tends to favor higher rates of sex than in non-subdivided populations, but that this effect often stays moderate.

## MODEL

### Life cycle

Our analytical model represents a diploid, facultatively sexual population structured according to the infinite island model, with discrete, non-overlapping generations (see Table 1 for an overview of the model’s parameters). At the start of each generation, *N* adults are present within each deme. Individual *j* in deme *i* produces a number of juveniles that is proportional to its fecundity *f*_*ij*_, which corresponds to the total amount of resources available for reproduction, and depends on the genotype of the individual (as described below). The investment in sex of the same individual is given by *σ*_*ij*_, and corresponds to the proportion of resources allocated to the production of female and male gametes, among all resources available for reproduction (the model assumes hermaphroditic individuals). The number of juveniles produced asexually by individual *j* in deme *i* is thus proportional to *f*_*ij*_ (1 − *σ*_*ij*_), while the number of juveniles produced sexually through female gametes is proportional to *f*_*ij*_ *σ*_*ij*_*/c*, where *c* represents the cost of sex (which may stem from resources invested in the production of male gametes, when those male gametes do not bring any resource to the fertilized egg): *c* = 1 in the absence of cost of sex, while *c* = 2 corresponds to a twofold cost of sex. The relative contribution of individual *j* to the pool of male gametes is proportional to *f*_*ij*_ *σ*_*ij*_. Female and male gametes are assumed to fuse at random within each deme. The total number of juveniles produced is assumed to be very large (effectively infinite). Asexual reproduction is equivalent to clonal reproduction, so that parent and offspring are genetically identical except for new mutations. Each juvenile disperses to another deme with a probability *d*; for simplicity, we assume that all demes contribute equally to the migrant pool (soft selection), so that the proportion of immigrant individuals in each deme after dispersal is given by *d*. Finally, *N* juveniles are randomly sampled within each deme to form the next adult generation (drift within demes).

**Table 1:** Parameters of the model.

|  |  |
| --- | --- |
| $N$ | Number of individuals per deme |
| $d$ | Dispersal rate |
| $s$ | Strength of selection against deleterious alleles $a, b$ |
| $h$ | Dominance coefficient of deleterious alleles |
| $u$ | Per-locus mutation rate towards deleterious alleles |
| $U$ | Deleterious mutation rate per haploid genome |
| $\sigma$ | Baseline rate of sex |
| $\delta\sigma$ | Effect of the modifier allele $m$ on the rate of sex |
| $h_m$ | Dominance coefficient of modifier allele $m$ |
| $c$ | Cost of sex (no cost of sex when $c = 1$) |
| $\sigma_e$ | Effective rate of sex $\sigma / [c(1 - \sigma) + \sigma]$ |
| $r_{ma}, r_{mb}, r_{ab}$ | Recombination rates between pairs of loci |
| $R$ | Chromosome map length |
| $\mu$ | Mutation rate at the modifier locus (in simulations) |
| $n_d$ | Number of demes (in simulations) |

### Genetic architecture

The investment in sex *σ*_*ij*_ of an individual depends on its genotype at a modifier locus. Two alleles *M* and *m* segregate at this locus, *σ*_*ij*_ being given by *σ, σ* + *δσh*_*m*_ and *σ* + *δσ* in *MM, Mm* and *mm* individuals, respectively (*δσ* thus represents the modifier effect, while *h*_*m*_ is the dominance coefficient of allele *m*). In the two-locus model, fertility *f*_*ij*_ is affected by a single selected locus at which a deleterious allele *a* is produced by mutation from the wild-type allele *A* at a rate *u* per generation. The fecundities of *AA, Aa* and *aa* individuals are proportional to 1, 1 − *sh* and 1 − *s*, respectively. In the three-locus model, fertility is also affected by a second selected locus with two alleles *B* and *b*, where allele *b* has the same selection (*s*) and dominance (*h*) coefficients as allele *a* at the first locus, and where mutation from *B* to *b* also occurs at a rate *u* per generation. We assume multiplicative effects of the two selected loci on fertility (no epistasis). To simplify the notation, *m, a* and *b* will also be used to identify the three loci, and *r*_*ma*_, *r*_*mb*_ and *r*_*ab*_ will denote the recombination rates between the different pairs of loci. As explained in File S3, our results can be extrapolated to the case of deleterious mutations occurring at a large number of loci along a linear chromosome, with a deleterious mutation rate *U* per chromosome per generation. While this involves integrating numerically a complicated expression over all possible positions of deleterious alleles along the chromosome (which is computationally demanding), the results are often very similar to those obtained when assuming free recombination among all loci (at least when the genetic map length of the chromosome is sufficiently large), and we thus often used the latter approximation.

### Allele frequencies and genetic associations

We define *p*_*m*(*ij*1)_, *p*_*m*(*ij*2)_ as indicator variables that equal 1 if allele *m* is present on the first or second haplotype (respectively) of individual *j* in deme *i*, and 0 otherwise. The frequency of allele *m* within individual *j* of deme *i* is denoted *p*_*m*(*ij*)_ = (*p*_*m*(*ij*1)_ + *p*_*m*(*ij*2)_) */*2, while the frequency of *m* in deme *i* is denoted *p*_*m*(*i*)_ = E_*j*_ [*p*_*m*(*ij*)_], where E_*j*_ stands for the average of all individuals *j* of deme *i*. The frequency of *m* in the whole metapopulation is denoted *p*_*m*_ = E_*i*_ [*p*_*m*(*i*)_], where E_*i*_ is the average of all demes *i*. Similar variables are defined for alleles *a* and *b*.

Genetic associations are defined using the notation of Barton and Turelli (1991) and Kirkpatrick et al. (2002), extended to the infinite island model of population structure by Roze and Rousset (2008). For example, *D*_*ma*_ measures the linkage disequilibrium (LD) between alleles *m* and *a*, defined as:

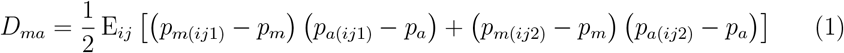

where E_*ij*_ stands for the average of all individuals *j* and all demes *i*. This is equivalent to *p*_*ma*_ − *p*_*m*_ *p*_*a*_, where *p*_*ma*_ is the frequency of *ma* chromosomes in the metapopulation. The association *D*_*a,a*_ is defined as:

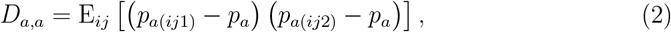

which is equivalent to *p*_*aa*_ − *p*_*a*_^2^ (where *p*_*aa*_ is the frequency of *aa* homozygotes in the metapopulation), thus measuring a deviation from Hardy-Weinberg equilibrium at the metapopulation level. This association can be expressed in terms of Wright’s *F*_IT_ statistic (Wright, 1951) as *D*_*a,a*_ = *F*_IT_ *p*_*a*_ *q*_*a*_ (where *q*_*a*_ = 1 − *p*_*a*_; Roze and Rousset, 2008). Similarly, *D*_*m,a*_ measures the association between alleles *m* and *a* present on different chromosomes of the same individual:

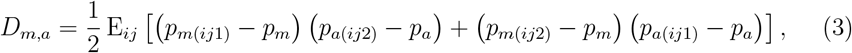

while the association *D*_*ma,a*_ is defined as:

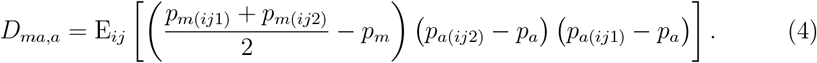

One can show that *D*_*ma,a*_ is positive when allele *m* is more frequent than allele *M* in *aa, AA* homozygotes, and negative when *m* is more frequent than *M* in *Aa* heterozygotes.

This notation can be extended to define associations between alleles present in two individuals sampled with replacement from the same deme. In particular,

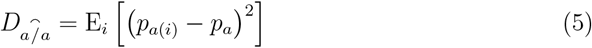

is the association between two alleles *a* in two individuals sampled with replacement from the same deme (which is also the variance in frequency of *a* among demes), expressed in terms of Wright’s *F*_ST_ statistic as *F*_ST_ *p*_*a*_ *q*_*a*_ (Wright, 1951; Roze and Rousset, 2008). Similarly, the associations 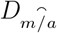 and 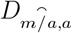 are defined as:

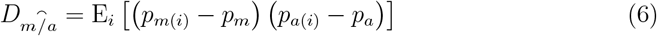

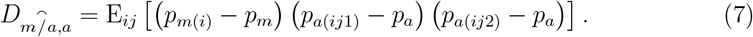

More general definitions, and a method to derive recurrence equations describing the changes in genetic associations from one generation to the next are given in File S1 (implemented in a *Mathematica* notebook available as Supplementary Material, https://doi.org/10.5281/zenodo.14670869), while Roze and Rousset (2008) should be consulted for further details on the general method.

### Quasi-equilibrium approximation

Once the frequencies of deleterious alleles *a* and *b* in the metapopulation have reached mutation-selection balance, all changes in allele frequencies (*p*_*m*_, *p*_*a*_, *p*_*b*_) are solely due to the modifier effect *δσ*, and are thus slow when *δσ* is small. Because genetic associations are broken by dispersal, sex and recombination, a separation of timescales argument can be used when *δσ* ≪ *d, σ, r*_*ij*_, as genetic associations will reach a quasi-equilibrium value on a relatively short timescale compared to the change in allele frequencies (Nagylaki, 1993; Barton and Turelli, 1991; Roze and Rousset, 2008). In this case, associations can be expressed in terms of allele frequencies and of the different parameters of the model. Throughout the paper, we will thus assume that *δσ* is small, and compute all expressions to the first order in *δσ*. Although the general method of Roze and Rousset (2008) can be used with arbitrary deme size *N* and dispersal rate *d*, it requires computing a large number of genetic associations in the two and three-locus diploid models when reproduction is partially clonal, leading to complicated expressions. In order to obtain tractable results, we will assume that deme size *N* is large, and compute all results to the first order in 1*/N* (this implicitly assumes that *Nd, Ns* ≫ 1, and therefore that the degree of population structure is weak). We also assume that the deleterious mutation rate per locus is small (*u* ≪ *s*) so that deleterious alleles stay at low frequency, and compute all results to leading order in *p*_*a*_, *p*_*b*_. Finally, we assume that selection against deleterious alleles is weak (*s* small but larger than 1*/N* and *u*, i.e., 1*/N, u* ≪ *s* ≪ 1), and compute recurrence equations to leading order in *s*.

### Simulations

Our analytical results were checked using simulation programs written in C++ and available from Zenodo (https://doi.org/10.5281/zenodo.14670869). One and two-locus simulations were used to verify analytical expressions for genetic associations at quasi-equilibrium in the two-locus model, and also to verify our expression for the linkage disequilibrium between deleterious alleles (*D*_*ab*_) in the three-locus model. These simulations represent a finite number of demes (*n*_d_), and keep track of the 3 (one-locus) or 10 (two-locus) diploid genotype frequencies within each deme. After 1000 preliminary generations, alleles frequencies and genetic associations are measured every 10 generations until obtaining a large number of measures (of the order 10^7^ – 10^8^). The frequencies of deleterious alleles (*p*_*a*_, *p*_*b*_) were initialized at zero; a rate of back mutation *v* = *u/*10 was introduced to prevent the permanent fixation of *a* or *b*. The frequency of the modifier allele *m* was initialized at *p*_*m*_ = 0.5; recurrent mutation was not introduced at the modifier locus, but the program was re-initialized when *p*_*m*_ reached values lower than 0.05 or higher than 0.95.

The multilocus, individual based model used in Roze and Michod (2010) was transposed to the case of a finite island model with *n*_d_ demes, to explore the evolution of the rate of sex when deleterious mutations occur at a very large number of possible sites along a linear chromosome, at a rate *U* per chromosome per generation. All mutations have the same selection and dominance coefficients (*s, h*) and have multiplicative effects on fitness. A sex modifier locus is located at the mid-point of the chromosome, controlling the investment in sex *σ* of the individual. An infinite number of possible alleles may occur at this locus, coding for different rates of sex between 0 and 1 (the investment in sex of an individual is given by the average of the values coded by its two alleles at the modifier locus). When reproduction is sexual, the number of crossovers at meiosis is drawn from a Poisson distribution with parameter *R* (chromosome map length in Morgans, set to 10 in order to mimic a genome with multiple chromosomes). During 1000 preliminary generations, the rate of sex is fixed to 1 in order to reach mutation-selection balance for deleterious alleles. Mutations are then introduced at the modifier locus at a rate *µ* = 10^*−*4^ per generation. When a mutation occurs, with probability 0.5 the new value of the modifier allele is drawn at random from a uniform distribution between 0 and 1, while with probability 0.5 the new value is drawn from a uniform distribution between *σ* − 0.1 and *σ* + 0.1, where *σ* is the value of the allele before mutation, in order to allow both small and large effect mutations (however, we observed that removing either type of mutation did not significantly affect equilibrium rates of sex). The simulations lasted for 2 × 10^6^ generations, the mean rate of sex, mean fitness and mean number of mutations per chromosome being measured every 100 generations. Besides the soft selection model (where all demes contribute equally to the migrant pool), we also considered a model in which the contribution of each deme to the migrant pool is proportional to the mean fitness of the deme; however, both models yielded very similar results for all parameter values tried (results not shown).

## RESULTS

### Two-locus model

We will focus here on the case where sex entails no direct cost (*c* = 1), and where alleles at the sex modifier locus have additive effects (*h*_*m*_ = 1*/*2); more general results for arbitrary *c* and *h*_*m*_ can be found in Files S1 and S2. In the absence of any direct cost of sex, the change in frequency of allele *m* is only due to genetic associations with the selected locus, and is given by:

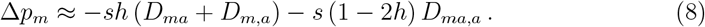

Associations *D*_*ma*_ and *D*_*m,a*_ are positive when allele *m* tends to be associated with the deleterious allele *a*, on the same (*D*_*ma*_) or on the homologous chromosome (*D*_*m,a*_), and negative when *m* tends to be associated with *A*. The association *D*_*ma,a*_ is positive when *m* tends to be more frequent among homozygotes at the selected locus, which is disadvantageous when *aa* and *AA* homozygotes have a lower fitness (on average) than *Aa* heterozygotes, that is, when *h <* 0.5. From File S2, an expression for *D*_*ma,a*_ at quasi-equilibrium is given by:

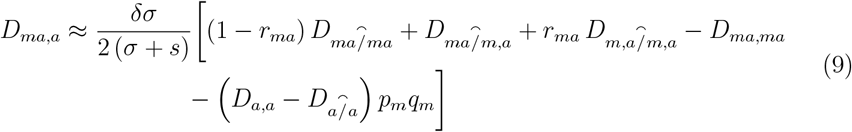

where *q*_*m*_ = 1 − *p*_*m*_. We will first discuss the term that appears on the second line of equation 9, and then the terms of the first line. Assuming that *p*_*a*_ and 1*/N* are small (so that only terms in *p*_*a*_^2^ and *p*_*a*_*/N* are retained), the single locus associations *D*_*a,a*_ and 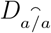 are given by (see File S1 for derivation):

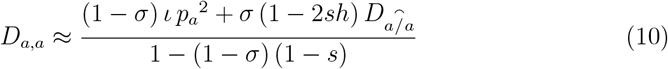

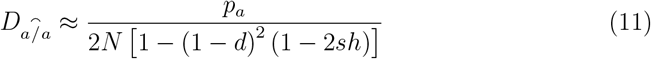

where *ι* = −*s* (1 − 2*h* + *h*^2^*s*) is a measure of dominance on a multiplicative scale (Otto, 2003). Given that *D*_*a,a*_ and 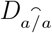 are equivalent to *F*_IT_ *p*_*a*_*q*_*a*_ and *F*_ST_ *p*_*a*_*q*_*a*_ (Roze and Rousset, 2008), the difference 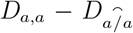 can be shown to be equivalent to *F*_IS_ *p*_*a*_*q*_*a*_, measuring an average deviation from Hardy-Weinberg equilibrium within demes (Wright, 1951). Indeed, *F*_IS_ is given by (*F*_IT_ − *F*_ST_) */* (1 − *F*_ST_), which is equivalent to 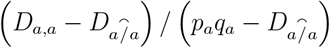 and thus to 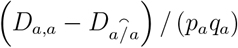 to leading order in *p*_*a*_ and 1*/N*. When *σ, s* and *d* are small (but still assuming 1*/N, p*_*a*_ ≪ *σ, s, d*), equations 10 – 11 yield:

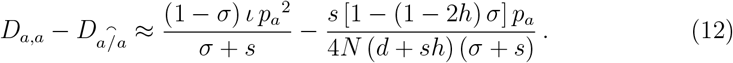

Equation 12 shows that two different effects generate a deviation from Hardy-Weinberg equilibrium within demes: dominance measured on a multiplicative scale (when ι ≠ 0, first term), and a combined effect of selection and population structure (second term). The first effect was already described by Otto (2003): in particular, when the deleterious allele *a* is partially recessive (*ι <* 0), selection generates an excess of heterozygotes which is maintained over generations when reproduction is at least partly clonal (*σ <* 1), leading to negative *D*_*a,a*_. The second term of equation 12 is always negative as long as *h >* 0, and is generated by selection and drift within demes.

Indeed, while both *D*_*a,a*_ and 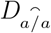 decrease during selection, 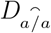 is increased by drift within demes while *D*_*a,a*_ stays unchanged, causing *D*_*a,a*_ to be lower than 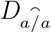 at the next generation (after drift). Figures 1 and 2 show that these results are confirmed by one-locus simulations. As can be seen in Figure 1, the first term of equation 12 (term in *ιp*_*a*_^2^) is negligible relative to the second term when the deleterious mutation rate *u* (and thus *p*_*a*_) is sufficiently small. Figure 2 shows that 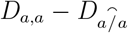 becomes more negative as *N, d* and *σ* decrease, our analytical approximation becoming less accurate when *N* is not large.

**Figure 1.**
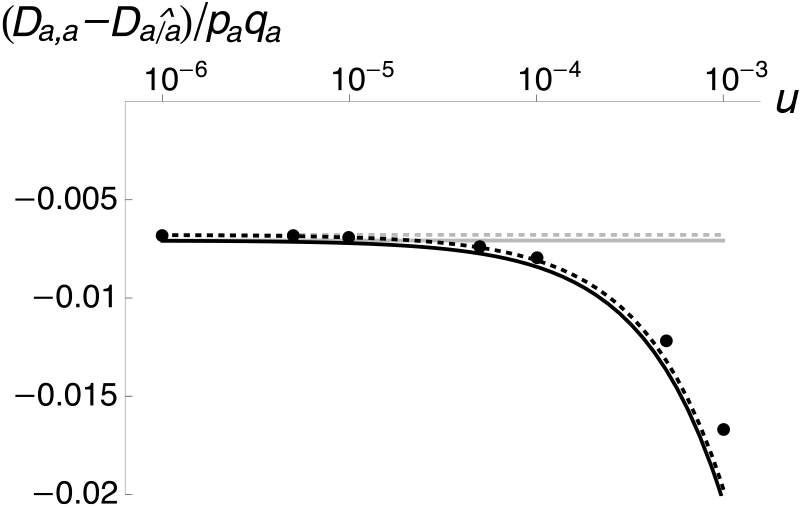
Average deviation from Hardy-Weinberg equilibrium within demes at a selected locus, measured as 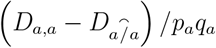 as a function of the rate of deleterious mutation *u* (on a log scale). The dots correspond to one-locus simulations results (error bars are smaller than the size of symbols), the black solid curve to the result obtained from equations 10 and 11, and the black dotted curve to equation 12. In these expressions, we used *p*_*a*_*q*_*a*_ ≈ *p*_*a*_ ≈ *u/* (*sh*). Grey lines correspond to the same predictions, neglecting the terms in *ι*. Values of the other parameters are *σ* = 0.1, *c* = 1, *s* = 0.04, *h* = 0.25, *N* = 500, *d* = 0.01 and *n*_d_ = 200 (in the simulations).

**Figure 2.**
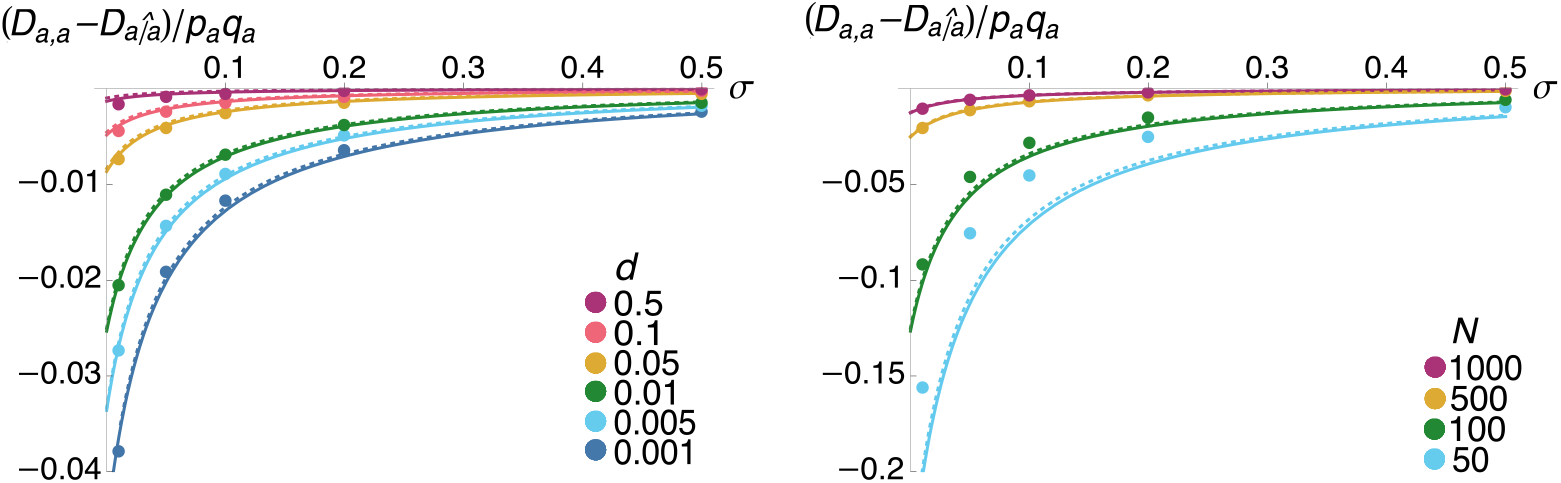
Average deviation from Hardy-Weinberg equilibrium within demes at a selected locus as a function of the rate of sex *σ*, for different values of the dispersal rate *d* (left figure) and deme size *N* (right figure). The dots correspond to one-locus simulation results (error bars are smaller than the size of symbols), solid curves to the result obtained from equations 10 and 11 and dotted curves to equation 12. Terms in *ι* have been neglected in these equations, but incorporating these terms leads to undistinguishable results. Parameter values are as in Figure 1, with *u* = 10^*−*5^.

Equation 12 also shows that *F*_IS_ = 0 at equilibrium in the absence of selection. This stands in contrast to the case of a single finite population, where an excess of heterozygotes is expected (on average) even in the absence of selection (Roze and Michod, 2010; Hartfield et al., 2016). This result can be recovered from our model in the absence of dispersal (*d* = 0): indeed, setting *s* = *d* = 0 in equations B1 and B2 of File S2, and subtracting B2 from B1 to obtain a recursion on 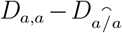 yields 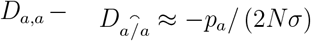 at equilibrium, in agreement with equation 2 in Roze and Michod (2010). Therefore, in the neutral case (*s* = 0), an excess of heterozygotes within demes is expected in the absence of dispersal (*d* = 0), but this effect disappears as soon as *d >* 0. This is also consistent with the results obtained by Balloux et al. (2003) from neutral probabilities of identity by descent in a finite island model, taking into account the fact that the definition of *F*_IS_ used here differs from the one used in Balloux et al. (2003). Indeed, *F*_IS_ as defined here may be expressed as (*Q*_0_ − *Q*_1_) */* (1 − *Q*_1_) where *Q*_0_ is the probability of identity by descent between the two alleles from the same individual and *Q*_1_ the probability of identity by descent between two alleles sampled with replacement from the same deme, while *F*_IS_ in Balloux et al. (2003) is defined as 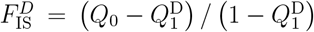 where 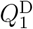 is the probability of identity by descent between two alleles sampled from two *different* individuals from the same deme (see also Rousset, 2004). Given that 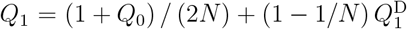 we have 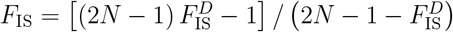 so that equation 12 in Balloux et al. (2003) yields *F*_IS_ = −1*/* [2*n*_d_ (*N* − 1) *σ* + 1] where *n*_d_ is the number of demes, showing that *F*_IS_ tends to zero as *n*_d_ tends to infinity, while *F*_IS_ is negative when *n*_d_ is finite.

As shown by equation 9, the excess of heterozygotes within demes 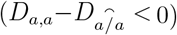 generates a positive *D*_*ma,a*_ association when allele *m* increases investment in sex (*δσ >* 0). Indeed, increasing the rate of sex decreases the proportion of heterozygotes among offspring, so that *m* tends to be more frequent than *M* among homozygotes at the selected locus. The first line of equation 9 shows that *D*_*ma,a*_ is also generated by several associations between the two loci. As explained in File S2, these associations stem from the variance in LD among demes, and the term on the first line of equation 9 is positive, also generating positive *D*_*ma,a*_ when *δσ >* 0. This comes from the fact that offspring produced by sexual reproduction within a deme (among which the frequency of *m* is higher when *δσ >* 0) tend to carry the allele at the selected locus with which *m* is associated in this deme (*a* is LD is locally positive, *A* if LD is locally negative), which also generates a positive association between *m* and homozygosity at the selected locus. An expression for *D*_*ma,a*_ at quasi-equilibrium can be obtained from the equations given in File S2 (see also *Mathematica* notebook available as Supplementary Material), and an approximation for small *s, d, σ* and *r*_*ma*_ is given by equation B35. When the deleterious allele *a* is partially recessive (*h <* 0.5), *D*_*ma,a*_ is positive when *δσ >* 0, so that the second term of equation 8 disfavors sex (due to the fact that sex produces more homozygous offspring, while homozygotes have a lower mean fitness than heterozygotes). When *a* is partially dominant (*h >* 0.5), *D*_*ma,a*_ is still positive when *δσ >* 0 as long as the deleterious mutation rate is sufficiently weak, so that the term in *ι* in equation 12 is negligible relative to the terms in 1*/N* generating positive *D*_*ma,a*_. In this case, the second term of equation 8 favors sex, since homozygotes have a higher mean fitness than heterozygotes.

From equations B37 – B38 in File S2, expressions for the pairwise associations *D*_*ma*_ and *D*_*m,a*_ at quasi-equilibrium are given by:

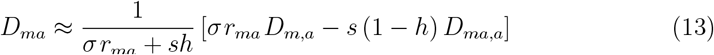

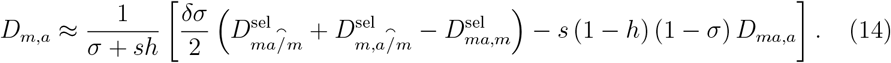

Equations 13 – 14 show that positive *D*_*ma,a*_ generates negative *D*_*ma*_ and *D*_*m,a*_. Indeed, when allele *m* is more frequent than *M* among homozygotes at the selected locus (*D*_*ma,a*_ *>* 0), selection against the deleterious allele *a* is more efficient among individuals carrying *m*, causing the frequency of *a* to become lower in those individuals (*D*_*ma*_, *D*_*m,a*_ *<* 0). This increased efficiency of selection against the deleterious allele favors higher rates of sex, through the first term of equation 8. Equation 14 shows that *D*_*ma*_ and *D*_*m,a*_ are also generated by the associations 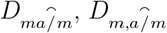 and *D*_*ma,m*_, whose values after selection appear in the first term of equation 14. As shown in File S2, these associations are generated by the variance in LD among demes and by selection against the deleterious allele, and are negative at quasi-equilibrium. Associations 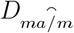 and 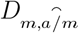 thus contribute to generate negative *D*_*ma*_ and *D*_*m,a*_, while the association *D*_*ma,m*_ has the opposite effect (see File S2 for biological interpretations of these associations and their effects). However, numerical analysis indicates that the effects of 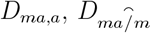 and 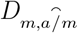 are always stronger than the effect of *D*_*ma,m*_, so that *D*_*ma*_ and *D*_*m,a*_ are negative at quasi-equilibrium, and favor higher rates of sex through the first term of equation 8.

When a direct cost of sex is introduced in the model (*c >* 1), the change in frequency of allele *m* can be decomposed into two elements:

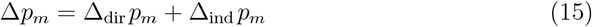

where Δ_dir_ *p*_*m*_ represents direct selection due to the cost of sex, and Δ_ind_ *p*_*m*_ indirect selection due to genetic associations with the selected locus. To leading order, the direct selection term is given by:

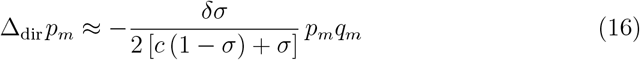

(see File S1). With a single selected locus, and assuming that the cost of sex is not very small, indirect selection is negligible compared with direct selection. However, when deleterious mutations occur at a large number of loci throughout the genome, the sum over all loci of terms in *p*_*a*_*/N* (that appear in Δ_ind_ *p*_*m*_) may become of the same order of magnitude as the direct selection term. As shown in File S2, the genetic associations involved in Δ_ind_ *p*_*m*_ are affected by the cost of sex. Indeed when *c >* 1, direct selection acts against both *m* and *a* and population structure generates interference effects between the two loci, which may change the sign of *D*_*ma*_, *D*_*m,a*_ and *D*_*ma,a*_ and generate new associations (see File S2). In this case, an expression for indirect selection is given by:

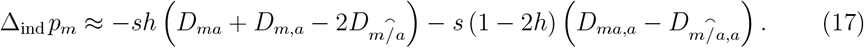

Figure 3 shows how the two terms of equation 17 vary with *σ*, once the associations that appear in equation 17 are replaced by their quasi-equilibrium values (obtained from the equations given in File S2, see also *Mathematica* notebook available as Supplementary Material). The dots in Figure 3 show the values obtained when genetic associations are measured directly from two-locus simulations (and averaged over a large number of points). These results show that our analytical approximations provide reasonable predictions of the values of genetic associations, and also show that adding a direct cost of sex may change the sign of the second term of equation 17.

**Figure 3.**
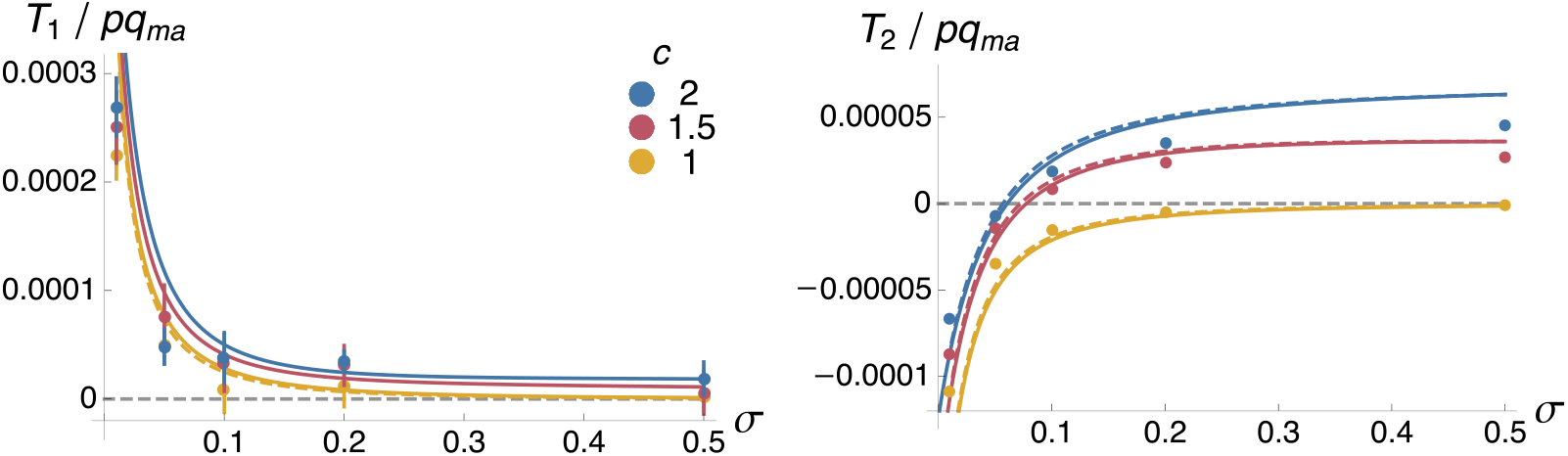
The two components of indirect selection for sex in the two-locus model, 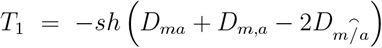 and 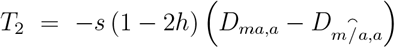 (from equation 17), as a function of the rate of sex *σ*. Both quantities are scaled by *pq*_*ma*_ = *p*_*m*_*q*_*m*_*p*_*a*_*q*_*a*_. The different colors correspond to different values of the cost of sex: *c* = 1 (yellow), 1.5 (red) and 2 (blue). Dots: two-locus simulation results. Solid curves: predictions obtained by solving the recurrence equations given in File S2 at equilibrium (terms in *ι* have been neglected from these equations). Dashed curve, left figure: prediction obtained using approximation B38–B39 in File S2 for *D*_*ma*_ + *D*_*m,a*_ when *c* = 1. Dashed curves, right figure: predictions obtained using approximation B28 in File S2 for *D*_*ma,a*_ (note that the association 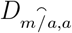 is always predicted to be negligible under our assumptions). Values of the other parameters are: *N* = 500, *d* = 0.01, *s* = 0.04, *h* = 0.25, *u* = 10^*−*5^, *δσ* = 0.02, *h*_*m*_ = 0.5, *r*_*ma*_ = 0.01 and *n*_d_ = 200 (in the simulations).

Figure 4 shows the evolutionarily stable (ES) rate of sex (towards which the population should evolve) predicted by extrapolations of the two locus model, when deleterious mutations occur at a very large number of loci (assuming all mutations have the same *s* and *h*). The dotted lines correspond to the results obtained when assuming free recombination among all loci, while the solid lines correspond to the case of a linear genome with a genetic map length of *R* = 10 Morgans. In the first case, the overall strength of indirect selection is obtained by replacing *r*_*ma*_ by 0.5 and *p*_*a*_ by *U/* (*sh*) in expressions of genetic associations at quasi-equilibrium, where *U* is the deleterious mutation rate per haploid genome. In the second case (linear genome), the overall strength of selection is obtained by expressing *r*_*ma*_ as a function of the genetic distance *x*_*ma*_ between loci using Haldane’s mapping function *r*_*ma*_ = [1 − exp (−2*x*_*ma*_)] */*2, and integrating over *x*_*ma*_ (see *Mathematica* notebook available as Supplementary Material). As can be seen on the figure, the results obtained under free recombination are very similar to those obtained with *R* = 10. When sex bears no direct cost (*c* = 1, Figure 4A), the two terms of equation 17 favor higher rates of sex when deleterious alleles are partially dominant (*h >* 0.5), so that the population is expected to evolve towards obligate sex (*σ* = 1). When deleterious alleles are partially recessive (*h <* 0.5), the second term of equation 17 disfavors sex (due to the immediate cost of increasing homozygosity among offspring, given that homozygotes have a lower mean fitness than heterozygotes when *h <* 0.5), while the first term of equation 17 favors sex (since sex increases the efficiency of selection against deleterious alleles), leading to an intermediate ES rate of sex. Similar results are obtained in the case of a single finite population (e.g., Figure 3 in Roze and Michod, 2010). Decreasing the dispersal rate *d* increases the ES rate of sex, but the effect stays relatively minor (indeed, decreasing *d* tends to increase the magnitude of both terms of equation 17). Population structure has a stronger effect on the ES rate of sex when sex is costly (*c* = 1.2, Figure 4B): in this case, the ES rate of sex results from a balance between direct and indirect selection, and the strength of indirect selection increases as *d* decreases, leading to higher ES rates of sex. Note that Figure 4 shows the effective rate of sex, defined as the proportion of sexually produced offspring, and given by *σ*_e_ = *σ/* [*c* (1 − *σ*) + *σ*] (*σ*_e_ = *σ* when *c* = 1).

**Figure 4.**
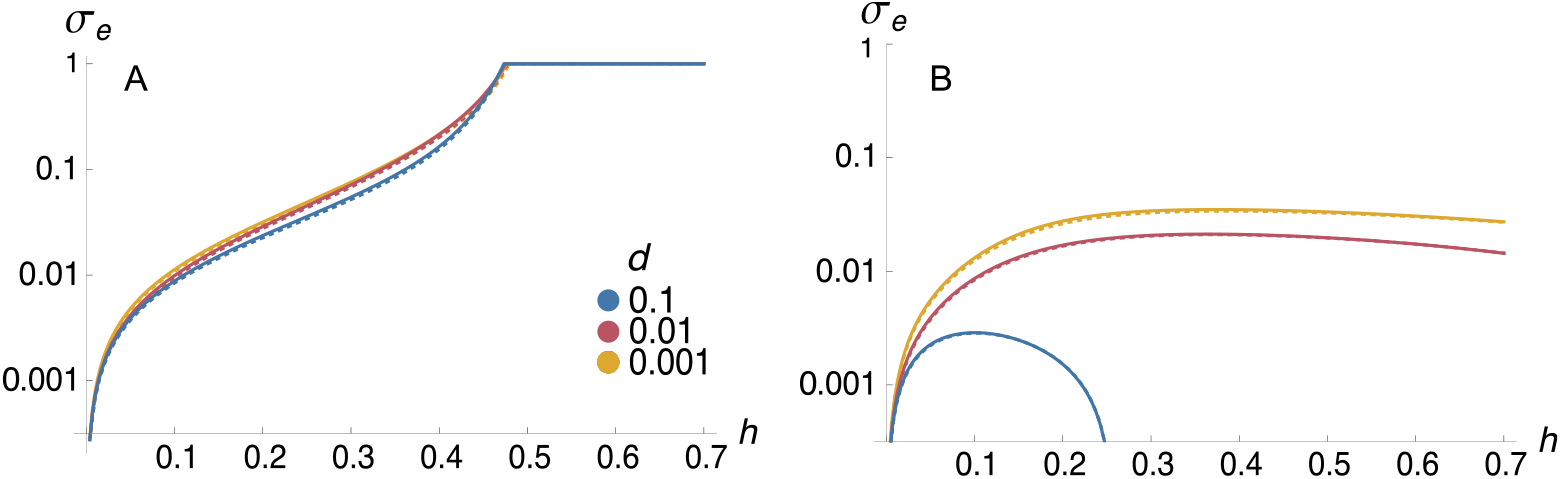
Evolutionarily stable (ES) effective rate of sex *σ*_e_ (on a log scale) predicted by extrapolating the two-locus model to the case of a whole genome with deleterious mutation rate *U*, as a function of the dominance coefficient *h* of deleterious alleles. The effective rate of sex corresponds to the proportion of sexually produced offspring, given by *σ*_e_ = *σ/*[*c*(1 − *σ*) + *σ*] (*σ*_*e*_ = *σ* when *c* = 1). The different colors correspond to different values of the dispersal rate *d*: *d* = 0.001 (yellow), 0.01 (red) and 0.1 (blue). Dotted curves correspond to the results obtained when assuming free recombination among all loci (*r*_*ma*_ = 0.5), while solid curves correspond to the case of a linear genome with a genetic map length of *R* = 10 Morgans (dotted and solid curves are nearly undistinguishable). A: *c* = 1 (no cost of sex); B: *c* = 1.2. Values of the other parameters are: *N* = 500, *U* = 0.2, *s* = 0.04.

### Three-locus model, and multilocus simulations

When deleterious mutations segregate at multiple loci, sex may benefit from additional advantages stemming from the effects of recombination between selected loci. In order to capture these effects, we extended our model to include a second selected locus, at which a deleterious allele *b* (with the same selection and dominance coefficients as allele *a* at the first selected locus) is segregating. The analysis of this three-locus model is presented in the Supplementary Material, and requires computing a relatively large number of genetic associations. We will not attempt here to provide a biological interpretation for each of these associations, but will focus on the linkage disequilibrium *D*_*ab*_ between deleterious alleles. The equations provided in File S3 show that *D*_*ab*_ is generated by several effects, and may be either negative or positive depending on parameter values. These effects ultimately stem from the variance in LD among demes caused by finite deme size, coupled with the effect of selection. Indeed, selection against deleterious alleles is more efficient when they are in positive LD than when they are in negative LD, so that *a* and *b* reach higher frequencies in demes where LD is negative than in demes where LD is positive. This tends to generate negative *D*_*ab*_ at the metapopulation level, corresponding to the classical Hill-Robertson effect (Hill and Robertson, 1966; Martin et al., 2006). However, the fact that *a* and *b* tend to be more frequent in demes where LD is negative generates a positive covariance between the local frequencies of deleterious alleles (measured by the association 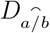), converted into positive *D*_*a,b*_ by gamete fusion, and into positive *D*_*ab*_ by recombination. Furthermore, the variance in LD among demes generates a positive covariance in homozygosity across loci, that also tends to generate positive *D*_*ab*_ — due to the fact that homozygosity increases the efficiency of selection against deleterious alleles (similarly, correlations in homozygosity across loci in partially selfing populations result in positive *D*_*ab*_, e.g., Stetsenko and Roze, 2022). Under obligate sexual reproduction (*σ* = 1), an approximation for *D*_*ab*_ when *d, s* and *r*_*ab*_ are small is given by:

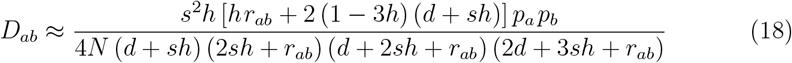

which is always positive when *h* ≤ 1*/*3, but may become negative when *h >* 1*/*3. Under complete dispersal (*d* = 1) and still for *σ* = 1, one obtains that *D*_*ab*_ should always be positive, and given by:

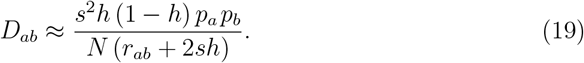

An approximation for *D*_*ab*_ when the rate of sex is low is given by equation C16 in File S3. Figure 5 compares the predictions obtained from these approximations (and from the more exact expressions obtained from the recurrence equations provided in File S3) with the results obtained from two-locus simulations representing the two selected loci. It illustrates that *D*_*ab*_ becomes more positive as *h* decreases, due to a stronger effect of the covariance in homozygosity among loci. The relative strength of the Hill-Robertson effect increases as the dispersal rate *d* decreases, *D*_*ab*_ becoming negative over wider ranges of values of *h. D*_*ab*_ is smaller in absolute value when *d* is large (Figure 5C), and is more often positive. In regimes where *D*_*ab*_ is negative under obligate sex (small *d, h* not too low), *D*_*ab*_ becomes more negative as the rate of sex *σ* decreases. By contrast, when *D*_*ab*_ is positive under obligate sex (higher *d*, lower *h*), *D*_*ab*_ often increases and then decreases as *σ* decreases. Note that *D*_*ab*_ represents the LD between deleterious alleles at the whole metapopulation level. As explained in File S3, the average LD within demes is given by 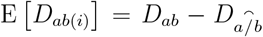 which, under obligate sex, is approximately:

**Figure 5.**
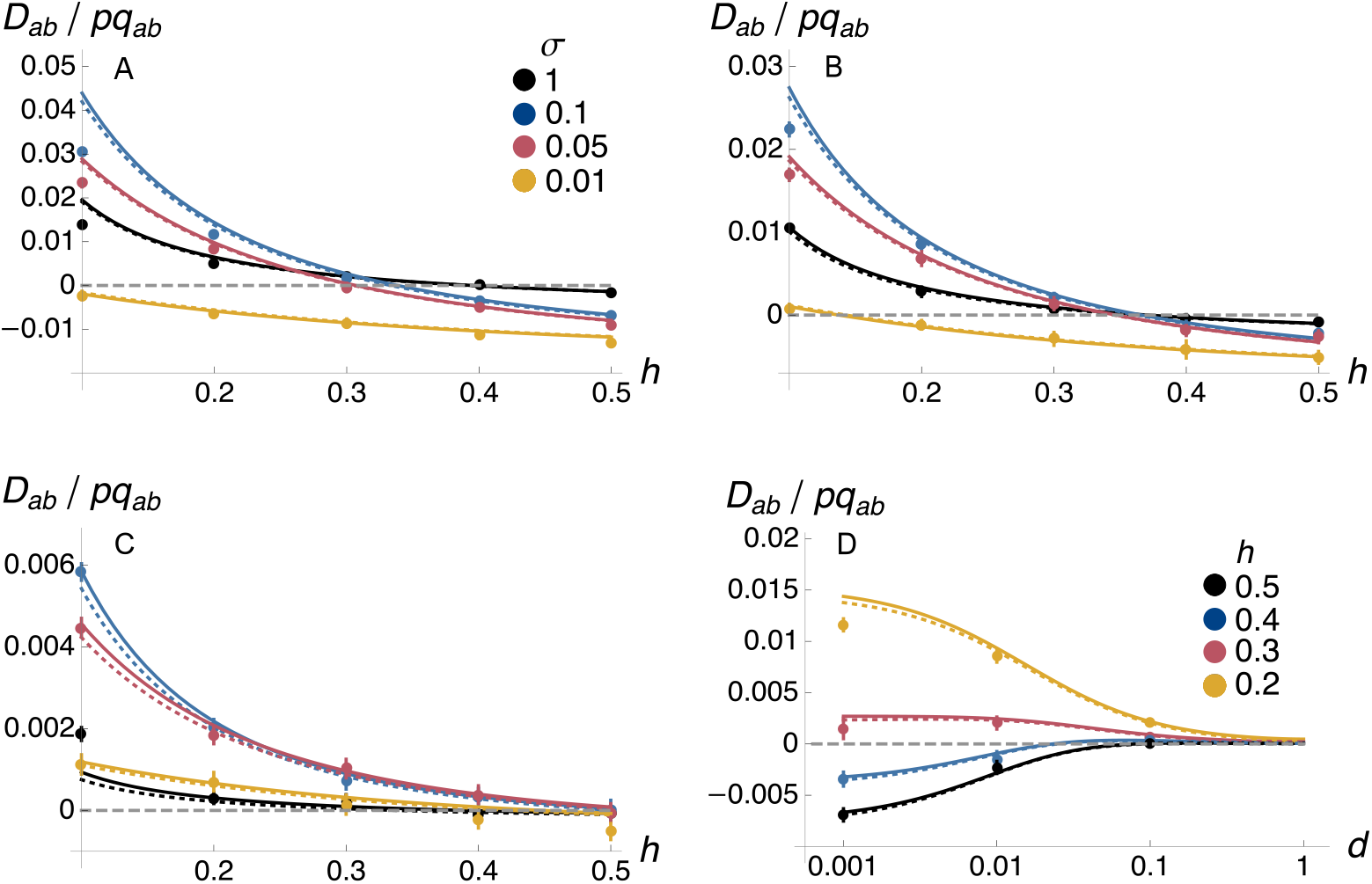
Linkage disequilibrium between deleterious alleles *a* and *b* (*D*_*ab*_, scaled by *pq*_*ab*_ = *p*_*a*_*q*_*a*_*p*_*b*_*q*_*b*_) as a function of the dominance coefficient of deleterious alleles *h* (A, B and C) and the dispersal rate *d* (D). The dots correspond to two-locus simulation results, solid curves to the predictions obtained from the recurrence equations given in File S3, and dotted curves to approximation 18 when *σ* = 1 and to equation C16 in File S3 when *σ <* 1. A: *d* = 0.001; B: *d* = 0.01; C: *d* = 0.1. In A, B, C, the different colors correspond to different values of the rate of sex *σ*: *σ* = 0.01 (yellow), 0.05 (red), 0.1 (blue) and 1 (black). In D, the rate of sex is set at *σ* = 0.1 and the different colors correspond to different values of the dominance coefficient of deleterious alleles *h*: *h* = 0.2 (yellow), 0.3 (red), 0.4 (blue) and 0.5 (black). In all figures, the selection coefficient *s* is set so that *sh* = 0.01 (i.e., *s* = 0.1, 0.05, 0.033, 0.025 and 0.02 when *h* = 0.1, 0.2, 0.3, 0.4 and 0.5, respectively). Values of the other parameters are: *N* = 500, *r*_*ab*_ = 0.01, *u* = 10^*−*4^ and *n*_d_ = 200 (in the simulations).

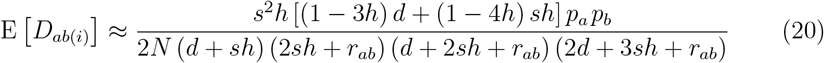

when *d, s* and *r*_*ab*_ are small. When the dispersal rate *d* is set to zero, equation 20 is equivalent to the result obtained by Roze (2021) in the case of a single finite population.

While recombination is beneficial in the presence of negative LD among deleterious alleles (as recombination increases the variance in fitness and the efficiency of selection), it should be disadvantageous when LD among deleterious alleles is positive.

Nevertheless, Roze (2021) showed that indirect selection on a recombination modifier in a finite diploid population is generated by many effects involving different genetic moments (covariances among genetic associations and allele frequencies), and that increased recombination rates are generally favored even in conditions leading to positive LD among deleterious alleles. Similarly in the present three-locus model, indirect selection at the sex modifier locus involves a large number of genetic associations among the three loci, generated by population structure, selection against deleterious alleles and the modifier effect (computed in the *Mathematica* notebook available as Supplementary Material). Providing an intuitive explanation for each of these associations is challenging, and their expressions at quasi-equilibrium generally take complicated forms. However, the model can be analyzed numerically and compared with the results obtained from the two-locus model, to assess the relative strength and direction of selection for sex generated by interactions between pairs of selected loci. As shown by Figure 6, the evolutionarily stable rate of sex predicted by extrapolating this three-locus model to the case of a whole genome with a deleterious mutation rate *U* (and assuming free recombination among all loci) is generally much higher that the prediction obtained from the two-locus model (compare solid and dotted lines on Figure 6). This indicates that interactions between selected loci make an important contribution to indirect selection, favoring higher rates of sex in most cases (except when *c* = 1 and *d* is large in Figure 6C).

**Figure 6.**
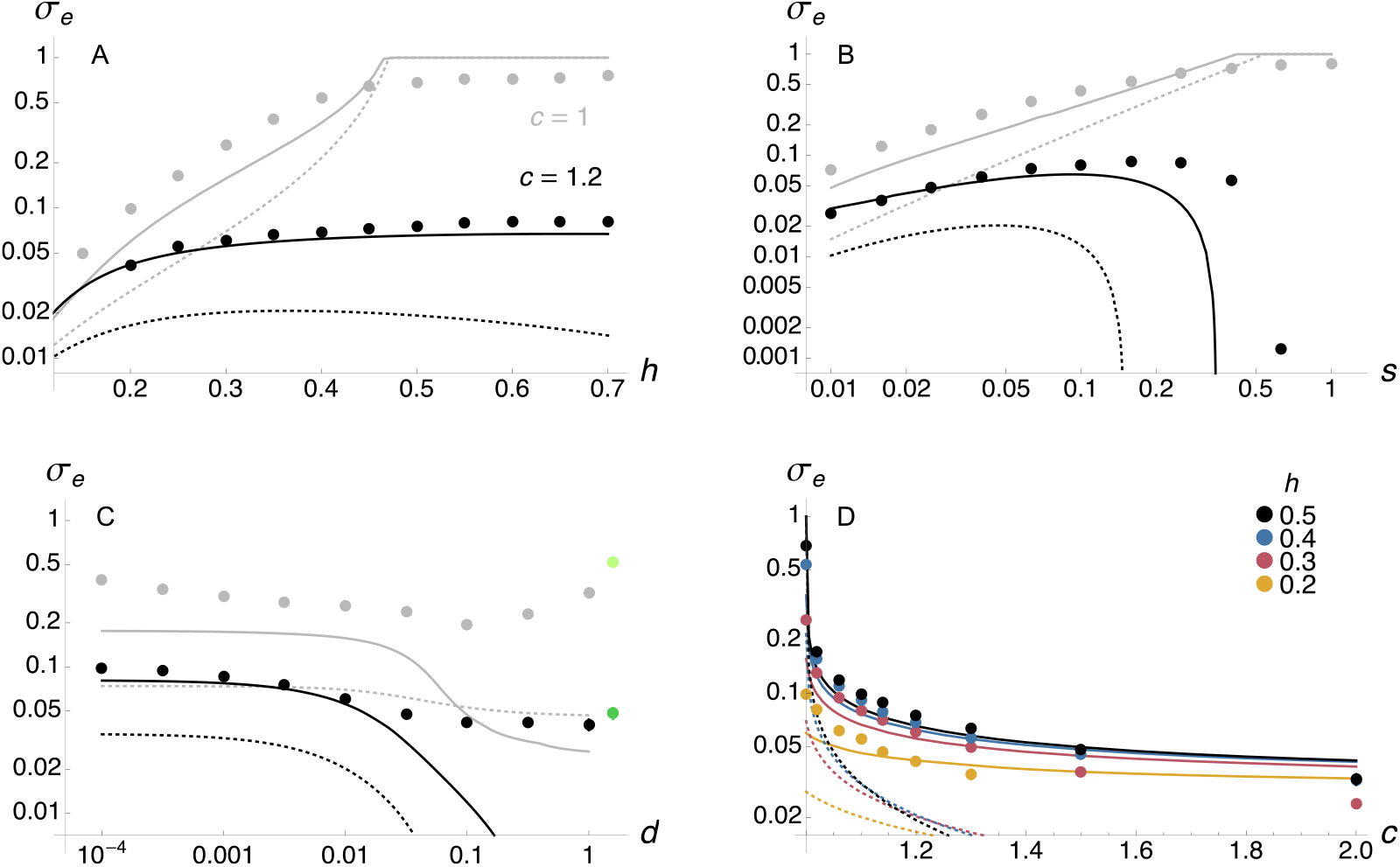
Evolutionarily stable effective rate of sex *σ*_*e*_ (on a log scale) when deleterious mutations occur at a rate *U* per haploid genome per generation, as a function of the dominance coefficient of deleterious alleles *h* (A), strength of selection against deleterious alleles *s* (B), dispersal rate *d* (C) and cost of sex *c* (D). Dots correspond to the average effective rate of sex at equilibrium in multilocus simulations (error bars are smaller than the size of symbols), solid curves to the predictions obtained from the three-locus model, and dotted curves to the predictions from the two-locus model (extrapolated to a whole genome, and assuming free recombination among all loci). Grey dots and curves in A, B, C correspond to the case where sex entails no direct cost (*c* = 1), while black dots and curves correspond to *c* = 1.2. Green dots on the right of Figure C correspond to the average effective rate of sex at equilibrium in simulations without any spatial structure (single population of size 10^5^), for *c* = 1 (lighter green) and *c* = 1.2 (darker green). In D, the different colors correspond to different values of *h*: *h* = 0.2 (yellow), 0.3 (red), 0.4 (blue) and 0.5 (black). Values of the other parameters are: *N* = 500, *d* = 0.01, *s* = 0.04, *h* = 0.3, *U* = 0.2. In the simulations the number of demes is set to *n*_d_ = 200, and the genome map length to *R* = 10 Morgans.

Comparisons with multilocus, individual-based simulation results (dots on Figure 6) show that extrapolations from our three-locus model often provide correct qualitative predictions of the effects of the different parameters on the ES rate of sex, but that quantitative (and sometimes qualitative) differences may arise. In particular, simulations confirm that the equilibrium rate of sex increases as the dominance coefficient of deleterious alleles *h* increases (Figure 6A; note that we could not obtain simulation results for values of *h* lower than the left-most dots in Figure 6A, as deleterious alleles were accumulating in the population). Increasing the strength of selection against deleterious alleles *s* leads to higher rates of sex when *c* = 1, while it has a non-monotonic effect when sex is costly (*c* = 1.2, Figure 6B). This is due to the fact that, although indirect selection favors higher rates of sex as *s* increases, the overall strength of indirect selection decreases when *s* becomes large, since deleterious alleles are rapidly eliminated from the population and genetic associations do no persist for long. Simulations show that the analytical model underestimates the value of *s* that maximizes the ES rate of sex, probably due to the fact that the model assumes weak selection. The simulations also confirm that decreasing the dispersal rate *d* only slightly increases the ES rate of sex when *d* is small (left part of Figure 6C). Stronger discrepancies between the analytical and simulation results occur for higher values of *d* (right part of Figure 6C), as the model predicts a strong decrease in the ES rate of sex as *d* increases, which is not observed in the simulations. This may be due to the fact that the finite size of the simulated metapopulation generates substantial indirect selection for sex even when the degree of population structure is weak (*d* high), preventing *σ* to reach very low values (by contrast, the analytical model assumes that the total population size is infinite). Indeed, the equilibrium rate of sex observed in the simulations when *d* is high is similar to its value in the case of a non-structured population with the same total size (green dots on Figure 6C). Therefore, when sex is costly (black dots in Figure 6C), increasing the degree of population structure (by decreasing the dispersal rate *d*) only leads to a moderate increase of the ES rate of sex.

When sex bears no direct cost (*c* = 1), the average rate of sex in the simulations is often larger than the value predicted by the analytical model, except when the model predicts obligate sex (for high values of *h* or *s*, Figures 6A and B), in which case lower rates of sex are observed in the simulations. This difference may be partly due to the fact that the analytical predictions assume free recombination among all loci, while the simulation program considers a linear chromosome (with a map length of 10 Morgans). Integrating the result of the three-locus model over a linear genome is computationally demanding, and was thus performed only for a few parameter values; however, the results obtained were only slightly higher than under free recombination (results not shown). A perhaps more likely explanation relies on the fact that indirect selection for sex is typically rather weak when *σ* is not small, so that mutation and drift at the modifier locus tend to displace the average rate of sex towards 0.5, corresponding to its average value under mutation and drift alone. Indeed, Figure S1 shows that decreasing the mutation rate at the modifier locus in the simulations (from *µ* = 10^*−*4^ to 10^*−*5^) reduces the discrepancy between the simulation and analytical results when *c* = 1. By contrast, decreasing *µ* does not have any noticeable effect on the average rate of sex when *c* = 1.2 (see Figure S1): in this case, stabilizing selection around the ES rate of sex is stronger, and the effect of mutational bias at the modifier locus is thus reduced. This is due to the fact that indirect selection for sex increases rapidly as *σ* decreases, so that sex is maintained in the population even when the cost of sex is strong, as shown by Figure 6D. As found by Roze and Michod (2010) in the case of a single finite population, we observed that when the cost of sex is high and *h* is sufficiently low, the rate of sex evolves to an intermediate value in the simulations, but at some point a mutant coding for a very low rate of sex invades and deleterious mutations then accumulate in the heterozygous state (preventing sex to increase again, as sexually produced offspring should be homozygous for many deleterious alleles and thus suffer from a strong segregation load). In the case of Figure 6D, this occurred for *h* = 0.2 and *c* = 1.5 and 2. Interestingly, population structure makes this invasion by asexual mutants less likely. This is shown in Figure 7 in the case of a twofold cost of sex (*c* = 2): in particular, for *h* = 0.3 the population becomes fully asexual when *d* ≥ 0.1, while an intermediate rate of sex is stably maintained over the 2 × 10^6^ generations of the simulation when *d <* 0.1. For *h* = 0.2, an intermediate rate of sex is stably maintained only for the lowest dispersal rate considered (*d* = 10^*−*4^). A possible explanation for these results is that population structure slows the spread of asexual mutants (Peck et al., 1999; Salathé et al., 2006; Hartfield et al., 2012): when a mutant coding for a very low (or zero) rate of sex occurs on a relatively fit genetic background, it tends to rapidly increase in frequency (due to the cost of sex); however its time to fixation is increased by population structure, giving more time for its relative fitness to decrease due to mutation accumulation.

**Figure 7.**
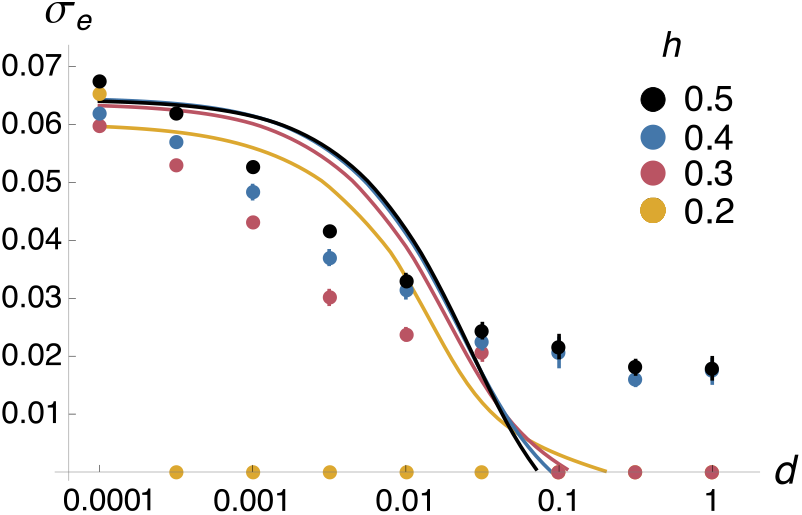
Evolutionarily stable effective rate of sex *σ*_*e*_ when deleterious mutations occur at a rate *U* per haploid genome per generation, as a function of the dispersal rate (*d*) and under a twofold cost of sex (*c* = 2). The different colors correspond to different values of *h*: *h* = 0.2 (yellow), 0.3 (red), 0.4 (blue) and 0.5 (black). Dots correspond to the average effective rate of sex at equilibrium in multilocus simulations (error bars are smaller than the size of symbols) and solid curves to the predictions obtained from the three-locus model (assuming free recombination among all loci). Values of the other parameters are as in Figure 6.

## DISCUSSION

The main genetic effect of sex is to mix genomes, and previous analytical models have explored how this may generate an advantage for sexual reproduction. In infinite populations without spatial structure, indirect selection for sex results from its effect in breaking genetic associations between alleles affecting fitness, either at the same locus or at different loci. In particular, dominance interactions lead to an excess or deficit of heterozygotes at selected loci (deviation from Hardy-Weinberg equilibrium), while epistatic interactions generate linkage disequilibrium (LD) between loci (Barton, 1995a; Otto, 2003). Additional sources of genetic associations stem from finite population size (Otto, 2021): in particular, genetic drift tends to generate an excess of heterozygotes at neutral and selected loci (Roze and Michod, 2010; Hartfield et al., 2016), while drift and selection generate LD between alleles affecting fitness, which may be either negative or positive depending on their degree of dominance (Hill and Robertson, 1966; Felsenstein, 1974; Roze, 2021). Furthermore, finite population models showed that indirect selection acting on a sex or recombination modifier is also affected by additional effects resulting from the variance in LD between the modifier and selected loci (Barton and Otto, 2005; Roze and Michod, 2010; Roze, 2014, 2021). Our model shows that similar effects arise in the case of a population subdivided into a very large number of finite demes, indirect selection for sex resulting from the interplay between a large number of mechanisms, corresponding to different sources of genetic associations between alleles (either in the same or in different individuals from the same deme). While these mechanisms can be understood individually in the case of a single selected locus, providing a complete heuristic description of the multiple effects resulting from interactions between selected loci is more challenging. Still, our analytical and simulation results lead to several conclusions on the effects of population structure on selection for sex.

First, our results show how population structure affects genetic associations that contribute to indirect selection for sex: in particular, deviations from Hardy-Weinberg equilibrium at selected loci, and LD between deleterious alleles. While drift alone tends to generate an excess of heterozygotes in isolated populations (particularly when the rate of sex is low), deviations from HWE within demes in a subdivided population are also affected by dispersal: in the island model and in the absence of selection, *F*_IS_ tends to zero on average when the number of demes tends to infinity (Balloux et al., 2003). Nevertheless, we have seen that selection against deleterious alleles and population structure combine to generate a local excess of heterozygotes at selected loci (negative *F*_IS_), for both partially recessive and partially dominant deleterious alleles. For realistic values of the deleterious mutation rate per locus, this effect is often much stronger than deviations from HWE generated by the deterministic effect of selection (Otto, 2003). By decreasing this local excess of heterozygotes, sex increases the variance in fitness and the efficiency of selection against deleterious alleles. When selection occurs at multiple loci, and in the regime considered here in which deleterious mutations stay near mutation-selection balance, interference between selected loci in an isolated, obligately sexual population generates negative LD between deleterious alleles when their dominance coefficient *h* is higher than 0.25, but positive LD when *h* is lower than 0.25 (Roze, 2021). Our results show that population structure tends to increase the parameter range in which LD is positive, in particular when the dispersal rate is high. This effect stems from correlations in homozygosity across loci (identity disequilibrium), resulting from the variance in the degree of inbreeding of individuals (more inbred individuals tend to be more homozygous, which increases the efficiency of selection at all loci, e.g.,Roze, 2009; Stetsenko and Roze, 2022). In the absence of epistasis, positive LD tends to disfavor sex, as recombination reduces the variance in fitness and the efficiency of selection.

A second result from our analytical model is that, in the absence of any direct cost of sex and when deleterious alleles are partially recessive, indirect selection should favor the maintenance of an intermediate rate of sex. Interactions between selected loci tend to increase substantially the evolutionarily stable rate of sex, even for parameter values leading to positive LD between deleterious alleles. Indeed (and as mentioned above), indirect selection for sex results from many different genetic effects, and most of these effects do not involve LD between deleterious alleles. As in previous single-population models (Roze and Michod, 2010; Roze, 2014), we found that the magnitude of indirect selection for sex increases rapidly as the rate of sex decreases, so that an intermediate rate of sex is often maintained even under substantial direct costs of sex. However, our simulation results show that invasion of asexual mutants may occur and prevent sex to be restored when deleterious mutations are partially recessive, as rare sexual individuals may then suffer from strong inbreeding depression. While our analytical model predicts that the evolutionarily stable rate of sex should increase with the degree of population structure, the simulations indicate that this increase is generally moderate: strongly structured populations do not show much higher rates of sex than weakly structured ones. The irreversible spread of asexual mutants occurs less frequently as the degree of population structure increases, probably due to the fact that asexual lineages have more time to decline in fitness before having reached fixation in the whole metapopulation (Peck et al., 1999; Salathé et al., 2006; Hartfield et al., 2012). Nevertheless, this irreversible spread is still often observed when *h* = 0.2 and when the cost of sex is strong (Figure 7).

While our model only shows limited effects of population structure on the evolutionarily stable rate of sex, stronger effects may possibly arise in different (and perhaps more realistic) models. In particular, models with variable deme size would be worth investigating, as local fluctuations in population size may amplify the effects of local drift and lead to stronger genetic associations. Furthermore, populations may better resist the invasion of asexual mutants if deme size could decrease as a result of mutation accumulation, so that demes in which asexual mutants have reached high frequency could shrink in size. Introducing local extinctions and recolonizations would represent another interesting extension of our model. While this would also increase local drift effects, high extinction rates may possibly favor asexuality due to the fact that the offspring of a small number of sexually reproducing colonizers may suffer from inbreeding depression. Based on this idea, Haag and Ebert (2004) proposed that frequent extinctions and recolonization of populations living in marginal habitats may explain the patterns of “geographic parthenogenesis” observed in many species (i.e., the fact that asexual lineages often tend to occupy marginal habitats, Glesener and Tilman, 1978; Lynch, 1984; Bierzychudek, 1985; Hörandl, 2009). This could be investigated further by using models in which marginal populations have higher risks of extinction. The effects of variable deme size and extinctions/recolonizations on the strength of selection for sex would probably be difficult to quantify analytically, but could be explored by simulation.

Many organisms live in heterogeneous environments. The effects of spatially changing environments and local adaptation on the selection for sex and recombination have been explored in previous models, which generally considered two demes containing an infinite number of individuals (no drift), connected by dispersal (Pylkov et al., 1998; Lenormand and Otto, 2000; Agrawal, 2009). In particular, Lenormand and Otto (2000) showed that in contrast to the case of a single population, recombination may increase the mean fitness of offspring when loci involved in local adaptation interact epistatically, increasing the range of epistasis where recombination is favored.

Similarly, segregation may increase the mean fitness of offspring when locally beneficial alleles tend to be dominant (Agrawal, 2009). These previous models considered one or two selected loci, and it would be interesting to obtain more quantitative results on the possible strength of selection for sex under different forms of environmental variation and genetic architectures of local adaptation. The effects of the interplay between local adaptation and local drift could also be explored by using models with finite deme size.

From an empirical perspective, more knowledge is needed on natural populations of facultatively sexual organisms, in order to asses to what extent the frequency of sexual reproduction may correlate with the degree of population structure or environmental heterogeneity. However, obtaining precise estimates of the rate of sex within natural populations may often be difficult. Experimental evolution on organisms displaying genetic variation in their relative investment in sexual and asexual reproduction (such as monogonont rotifers, e.g., Becks and Agrawal, 2010, 2012) may represent another possible approach to explore how population structure may affect the evolutionary advantage of sex.

## Acknowledgements

We thank the bioinformatics and computing services at Roscoff’s Biological Station (Abims platform) and at Institute of Science and Technology Austria for computing time. L.F. acknowledges the support of the NOMIS-ISTA Fellowship Program. DR was financed by the Centre National de la Recherche Scientifique (CNRS).

