## Supplementary material for "Deleterious mutations and selection for sex in spatially structured, diploid populations": File S1

### FILE S1: DERIVING RECURSIONS ON ALLELE FREQUENCIES AND GENETIC ASSOCIATIONS

The method of Roze and Rousset (2008) can be used to obtain recurrence equations describing the changes in allele frequencies and genetic associations over the different steps of the life cycle. The method has been implemented in a *Mathematica* notebook (available as Supplementary Material) that can be used to generate recursions automatically. We outline here the general method and illustrate it with some examples; Roze and Rousset (2008) should be consulted for further details.

**Variables.** Genetic associations are defined using the notation of Barton and Turelli (1991) and Kirkpatrick et al. (2002), extended to the infinite island model of population structure by Roze and Rousset (2008). We define $p_{a(ij1)}$, $p_{a(ij2)}$ as indicator variables that equal 1 if allele $a$ is present on the first or second haplotype (respectively) of individual $j$ in deme $i$, and 0 otherwise. Similarly, $p_{m(ij1)}$, $p_{m(ij2)}$ equal 1 if allele $m$ is present on the first or second haplotype of individual $j$ in deme $i$ (and 0 otherwise). The frequency of allele $a$ within individual $j$ of deme $i$ is denoted $p_{a(ij)} = \left(p_{a(ij1)} + p_{a(ij2)}\right)/2$, while the frequency of $a$ in deme $i$ is denoted $p_{a(i)} = \mathrm{E}_j\left[p_{a(ij)}\right]$, where $\mathrm{E}_j$ stands for the average of all individuals $j$ of deme $i$. The frequency of $a$ in the whole metapopulation is denoted $p_a = \mathrm{E}_{ij}\left[p_{a(ij)}\right]$, where $\mathrm{E}_{ij}$ is the average of all individuals $j$ and over all demes $i$. Variables $p_{m(ij)}$, $p_{m(i)}$ and $p_m$ are defined similarly.

Centered variables are defined as $\zeta_{a(ij1)} = p_{a(ij1)} - p_a$, $\zeta_{a(ij2)} = p_{a(ij2)} - p_a$, $\zeta_{a(ij)} = p_{a(ij)} - p_a$ and $\zeta_{a(i)} = p_{a(i)} - p_a$ (and similarly for $\zeta_{m(ij1)}$, $\zeta_{m(ij2)}$, $\zeta_{m(ij)}$ and $\zeta_{m(i)}$). Genetic associations between alleles present at different loci and/or different haplotypes of an individual are then defined as:

$$D_{\mathbb{U},\mathbb{V}} = \mathrm{E}_{ij}\left[\zeta_{\mathbb{U},\mathbb{V}(ij)}\right] \tag{A1}$$

with

$$\zeta_{\mathbb{U},\mathbb{V}(ij)} = \frac{1}{2}\left(\prod_{k\in\mathbb{U}}\zeta_{k(ij1)}\prod_{l\in\mathbb{V}}\zeta_{l(ij2)} + \prod_{k\in\mathbb{U}}\zeta_{k(ij2)}\prod_{l\in\mathbb{V}}\zeta_{l(ij1)}\right) \tag{A2}$$

and where $\mathbb{U}$, $\mathbb{V}$ are sets that can contain any combination of lowercase alleles at both loci, namely $\varnothing$, $m$, $a$ and $ma$. For simplicity, associations between alleles on the same haplotype of an individual $D_{\mathbb{U},\varnothing} = D_{\varnothing,\mathbb{U}}$ will be denoted $D_{\mathbb{U}}$ — with two loci, this only concerns the linkage disequilibrium $D_{ma}$ between alleles $m$ and $a$. The within-locus association $D_{a,a}$ is a measure of excess homozygosity at the selected locus (it is equal to $p_{aa} - {p_a}^2$, where $p_{aa}$ is the frequency of $aa$ homozygotes in the whole metapopulation). $D_{m,a}$ measures the association between $m$ an $a$ on different haplotypes of the same individual, while $D_{ma,a}$ tends to be positive when allele $m$ is more frequent than $M$ among homozygotes at the selected locus ($aa$ or $AA$), and negative when $m$ is more frequent than $M$ among $Aa$ individuals.

This notation can be extended to define associations between alleles present in different individuals from the same deme, sampled either with or without replacement among all individuals from a deme (Roze and Rousset, 2008). In particular, $D_{\mathbb{U},\mathbb{V}\widehat{/}\mathbb{S},\mathbb{T}}$ stands for the association among sets of alleles $\mathbb{U}$ and $\mathbb{V}$ on different haplotypes of an individual, and sets of alleles $\mathbb{S}$ and $\mathbb{T}$ on different haplotypes of another individual, sampled with replacement from the same deme:

$$D_{\mathbb{U},\mathbb{V}\widehat{/}\mathbb{S},\mathbb{T}} = \mathrm{E}_{ijk}\left[\zeta_{\mathbb{U},\mathbb{V}(ij)}\,\zeta_{\mathbb{S},\mathbb{T}(ik)}\right] \tag{A3}$$

where $\mathrm{E}_{ijk}$ stands for the average over all demes $i$ and over all pairs of individuals $j$, $k$ (including $j = k$). Finally, $D_{\mathbb{U},\mathbb{V}/\mathbb{S},\mathbb{T}}$ stands for the same association when individuals are sampled without replacement from a deme:

$$D_{\mathbb{U},\mathbb{V}/\mathbb{S},\mathbb{T}} = \mathrm{E}_{ij,k\neq j}\left[\zeta_{\mathbb{U},\mathbb{V}(ij)}\,\zeta_{\mathbb{S},\mathbb{T}(ik)}\right] \tag{A4}$$

where $\mathrm{E}_{ij,k\neq j}$ stands for the average over all demes $i$ and over all pairs of individuals $j$, $k$ with $j \neq k$. Note that we have:

$$D_{\mathbb{U},\mathbb{V}\widehat{/}\mathbb{S},\mathbb{T}} = \frac{1}{2N}\left(D_{\mathbb{US},\mathbb{VT}} + D_{\mathbb{UT},\mathbb{VS}}\right) + \left(1 - \frac{1}{N}\right) D_{\mathbb{U},\mathbb{V}/\mathbb{S},\mathbb{T}}, \tag{A5}$$

as two individuals sampled with replacement from a deme are the same individual with probability $1/N$. For example $D_{a\widehat{/}a} = \mathrm{E}_{ijk}\left[\zeta_{a(ij)}\,\zeta_{a(ik)}\right] = \mathrm{E}_i\left[{\zeta_{a(i)}}^2\right]$ is the association between two alleles $a$ at the selected locus among individuals sampled with replacement from the same deme, which is equivalent to the variance in the frequency of $a$ among demes (variance of $p_{a(i)}$).

When deriving recurrence equations on allele frequencies and genetic associations, it proves convenient to decompose the production of juveniles into two steps:

"selection" corresponds to the effect on allele frequencies and genetic associations of differences in fertilities ($f_{ij}$) among parents due to different genotypes at the selected locus, while "reproduction" corresponds to the effect of juvenile production through asexual and sexual reproduction, at rates that depend on the genotype of the parent at the modifier locus (this step includes the effects of meiosis and fertilization in the case of sexually produced offspring). Associations measured after selection, reproduction, dispersal and drift within demes will be denoted $D_{\mathcal{S}}^{\text{sel}}$, $D_{\mathcal{S}}^{\text{repr}}$, $D_{\mathcal{S}}^{\text{disp}}$ and $D_{\mathcal{S}}{}'$ (respectively) where $\mathcal{S}$ may be any set of alleles present in one or several individuals from the same deme. Note that allele frequencies in the whole metapopulation only change during selection (due to differences in fertilities) and during reproduction (due to the cost of sex); $p_a^{\text{sel}}$ and $p_a{}'$ will denote the frequency of allele $a$ in the metapopulation after selection and the next generation (respectively), and similarly for $p_m^{\text{sel}}$, $p_m{}'$. We assume that deme size $N$ is large (with $Nd$, $Ns \gg 1$), the modifier effect $\delta\sigma$ is small, the mutation rate $u$ towards the deleterious allele $a$ is small ($u \ll s$) and $h$ is not too close to zero, so that $p_a$ at mutation-selection balance is small and $\approx u/(hs)$, while the effect of mutation on genetic associations may be neglected. Throughout the following, only the terms of order $\delta\sigma\, p_a{}^2$ and $\delta\sigma\, p_a/N$ appearing in the change in frequency of the modifier will be retained (in particular, all terms proportional to $p_a{}^2/N$ or $p_a/N^2$ will be ignored).

**Change in frequency of the modifier.** The change in frequency of allele $m$ during selection is given by:

$$\Delta_{\text{sel}} p_m = \mathrm{E}_{ij}\left[\frac{f_{ij}}{f_i}\, p_{m(ij)}\right] - p_m \tag{A6}$$

where $f_i$ is the average fecundity in deme $i$. Using the fact that $\mathrm{E}_{ij}\left[f_{ij}/f_i\right] = 1$, this is:

$$\Delta_{\text{sel}} p_m = \mathrm{E}_{ij}\left[\frac{f_{ij}}{f_i}\, \zeta_{m(ij)}\right]. \tag{A7}$$

The fecundity of individual $j$ in deme $i$ can be written as:

$$f_{ij} = 1 - s\left[h\left(p_{a(ij1)} + p_{a(ij2)}\right) + (1-2h)\, p_{a(ij1)} p_{a(ij2)}\right] \tag{A8}$$

After rearranging, this gives:

$$\frac{f_{ij}}{f_i} = \frac{1 - s\,T_a - 2s\left[h + (1-2h)\, p_a\right]\zeta_{a(ij)} - s\,(1-2h)\,\zeta_{a,a(ij)}}{1 - s\,T_a - 2s\left[h + (1-2h)\, p_a\right]\zeta_{a(i)} - s\,(1-2h)\,\zeta_{a,a(i)}} \tag{A9}$$

with $T_a = 2h\,p_a + (1-2h)\,{p_a}^2$. To make progress, we can assume that $s$ is small and express equation A9 to the first order in $s$, yielding:

$$\frac{f_{ij}}{f_i} = 1 - 2sh\left(\zeta_{a(ij)} - \zeta_{a(i)}\right) - s\,(1-2h)\left(\zeta_{a,a(ij)} - \zeta_{a,a(i)}\right) + o(s) \qquad \text{(A10)}$$

where the term in $p_a$ has been neglected. Alternatively, equation A9 may be expressed to leading order in $p_a$ (without any assumption on $s$), using the fact that $\zeta_{a(i)}$ and $\zeta_{a,a(i)}$ are of order $p_a$ under our assumptions; this also leads to equation A10, with an extra term of order $s^2$:

$$-4s^2\left[h\,\zeta_{a(ij)} + (1-2h)\,\zeta_{a,a(ij)}\right]\left[h\,\zeta_{a(i)} + (1-2h)\,\zeta_{a,a(i)}\right]. \qquad \text{(A11)}$$

Equations A7 and A10 yield:

$$\Delta_{\text{sel}}p_m \approx -sh\left(D_{ma} + D_{m,a} - 2D_{m\widehat{/}a}\right) - s\,(1-2h)\left(D_{ma,a} - D_{m\widehat{/}a,a}\right). \qquad \text{(A12)}$$

The term shown in equation A11 generates an extra term $-2s^2h^2\left(D_{ma\widehat{/}a} + D_{m,a\widehat{/}a}\right)$, which should be negligible unless $s$ is large (the other associations generated by equation A11 can be shown to be of order $1/N^2$).

The frequency of allele $m$ also changes during reproduction, when the reproductive mode affects the number of juveniles produced (*i.e.*, when $c \neq 1$). This change is given by:

$$\begin{aligned}\Delta_{\text{repr}}p_m = {} & \mathrm{E}^{\text{sel}}_{ij}\left[\frac{1-\sigma_i}{1-\sigma_i+\sigma_i/c}\,\frac{1-\sigma_{ij}}{1-\sigma_i}\,p_{m(ij)}\right] \\ & + \mathrm{E}^{\text{sel}}_{ijk}\left[\frac{\sigma_i/c}{1-\sigma_i+\sigma_i/c}\,\frac{\sigma_{ij}\sigma_{ik}}{{\sigma_i}^2}\,\frac{p_{m(ij)}+p_{m(ik)}}{2}\right] - p^{\text{sel}}_m\end{aligned} \qquad \text{(A13)}$$

where $\mathrm{E}^{\text{sel}}_{ij}$ and $\mathrm{E}^{\text{sel}}_{ijk}$ are averages over parents after selection, and $p^{\text{sel}}_m$ the frequency of $m$ after selection (before reproduction), while $\sigma_i$ is the average investment in sex in deme $i$. The first term of equation A13 corresponds to the frequency of $m$ among asexually produced offspring: $(1-\sigma_i)\,/\,(1-\sigma_i+\sigma_i/c)$ is the proportion of asexually produced offspring in deme $i$, while $(1-\sigma_{ij})\,/\,(1-\sigma_i)$ is the relative contribution of parent $j$ to the pool of asexually produced offspring in deme $i$, and $p_{m(ij)}$ the frequency of $m$ within this parent. The second term of equation A13 corresponds to the frequency of $m$ among sexually produced offspring: $(\sigma_i/c)\,/\,(1-\sigma_i+\sigma_i/c)$ is the proportion of sexually produced offspring in deme $i$, while $\sigma_{ij}/\sigma_i$, $\sigma_{ik}/\sigma_i$ are the relative contributions of parents $j$ and $k$ to the pool of gametes, and $\left(p_{m(ij)} + p_{m(ik)}\right)/2$ the average frequency of $m$ within these parents. After rearranging, this gives:

$$\Delta_{\text{repr}} p_m = \mathrm{E}_{ij}^{\text{sel}} \left[ \frac{1 - \sigma_{ij}}{1 - \sigma_i + \sigma_i / c} \zeta_{m(ij)} \right] + \mathrm{E}_{ijk}^{\text{sel}} \left[ \frac{\sigma_{ij} \sigma_{ik}}{c\, \sigma_i \left[1 - \sigma_i + \sigma_i / c\right]} \frac{\zeta_{m(ij)} + \zeta_{m(ik)}}{2} \right]. \tag{A14}$$

The investment in sex of parent $j$ in deme $i$ can be expressed as:

$$\sigma_{ij} = \sigma + \delta\sigma \left[ h_m \left( p_{m(ij1)} + p_{m(ij2)} \right) + (1 - 2h_m)\, p_{m(ij1)} p_{m(ij2)} \right], \tag{A15}$$

yielding:

$$\sigma_{ij} = \overline{\sigma} + 2\delta\sigma_m\, \zeta_{m(ij)} + \delta\sigma_{m,m} \left( \zeta_{m,m(ij)} - D_{m,m}^{\text{sel}} \right) \tag{A16}$$

$$\sigma_i = \overline{\sigma} + 2\delta\sigma_m\, \zeta_{m(i)} + \delta\sigma_{m,m} \left( \zeta_{m,m(i)} - D_{m,m}^{\text{sel}} \right) \tag{A17}$$

with

$$\overline{\sigma} = \sigma + 2\delta\sigma h_m\, p_m^{\text{sel}} + \delta\sigma\, (1 - 2h_m) \left[ \left( p_m^{\text{sel}} \right)^2 + D_{m,m}^{\text{sel}} \right] \tag{A18}$$

$$\delta\sigma_m = \delta\sigma \left[ h_m + (1 - 2h_m)\, p_m^{\text{sel}} \right], \qquad \delta\sigma_{m,m} = \delta\sigma\, (1 - 2h_m). \tag{A19}$$

Expressing the fractions in equation A14 to the first order in $\delta\sigma$ leads to an expression for $\Delta_{\text{repr}} p_m$ in terms of associations $D_{mm}$, $D_{m,m}$, $D_{m\widehat{/}m}$, $D_{mm,m}$ and $D_{m,m\widehat{/}m}$ measured among parents after selection. Repeated indices in associations can be eliminated using equation 5 in Kirkpatrick et al. (2002):

$$D_{ll\mathcal{S}} = p_l q_l D_{\mathcal{S}} + (1 - 2p_l)\, D_{l\mathcal{S}} \tag{A20}$$

where $\mathcal{S}$ can be any set of alleles from one or several individuals from the same deme, and with $q_l = 1 - p_l$. In particular, we have $D_{mm} = p_m q_m$ and $D_{mm,m} = (1 - 2p_m)\, D_{m,m}$. Furthermore, the association $D_{m,m\widehat{/}m}$ can be shown to be of order $1/N^2$, which finally leads to:

$$\Delta_{\text{repr}} p_m = -\,(c - 1) \left[ \delta\sigma_{m,\text{e}} \left( p_m q_m + D_{m,m}^{\text{sel}} - 2 D_{m\widehat{/}m}^{\text{sel}} \right) + \delta\sigma_{m,m,\text{e}}\, (1 - 2p_m)\, D_{m,m}^{\text{sel}} \right] \tag{A21}$$

with

$$\delta\sigma_{m,\text{e}} = \frac{\delta\sigma_m}{c\,(1 - \sigma) + \sigma}, \qquad \delta\sigma_{m,m,\text{e}} = \frac{\delta\sigma_{m,m}}{c\,(1 - \sigma) + \sigma}. \tag{A22}$$

Note that although the value of $p_m q_m$ in equation A21 should be taken after selection, the change in $p_m$ during selection is of order $\delta\sigma$ (as it involves associations between the modifier and the selected locus — shown in equation A12 — which are generated by the modifier effect), and would thus generate a term of order $(\delta\sigma)^2$ in the expression of $\Delta_{\text{repr}} p_m$. Furthermore, because we assume that the degree of population structure is weak ($Nd \gg 1$), the associations $D_{m,m}$ and $D_{m\overset{\frown}{/}m}$ should remain small (of order $1/N$) and may thus be neglected relative to the term in $p_m q_m$ of equation A21, yielding $\Delta_{\text{repr}} p_m \approx -\,(c-1)\,\delta\sigma_{m,\text{e}}\, p_m q_m$.

In the following, we explain how recurrence equations for the associations $D_{ma,a}$, $D_{a,a}$ and $D_{a\overset{\frown}{/}a}$ are derived; recursions for other associations can be obtained using the same principles, and are computed automatically in the *Mathematica* notebook available as Supplementary Material.

**Recursion on $\boldsymbol{D_{ma,a}}$.** The association $D_{ma,a}$ remains unchanged during dispersal and drift within demes, since it involves alleles present in the same individual: therefore, $D'_{ma,a} = D^{\text{disp}}_{ma,a} = D^{\text{repr}}_{ma,a}$. Computing the effect of reproduction on $D_{ma,a}$ can be decomposed into two steps (Barton and Turelli, 1991; Kirkpatrick et al., 2002; Roze and Rousset, 2008). We have:

$$D^{\text{repr}}_{ma,a} = \mathrm{E}^{\text{repr}}_{ij}\left[\left(p_{m(ij)} - p'_m\right)\left(p_{a(ij1)} - p'_a\right)\left(p_{a(ij2)} - p'_a\right)\right] \tag{A23}$$

where $\mathrm{E}^{\text{repr}}_{ij}$ stands for the average over all demes and individuals after reproduction, and $p'_m$, $p'_a$ are allele frequencies after reproduction. This can also be written as:

$$\begin{aligned} D^{\text{repr}}_{ma,a} = \mathrm{E}^{\text{repr}}_{ij}\Big[&\left(p_{m(ij)} - p^{\text{sel}}_m - \Delta_{\text{repr}} p_m\right)\left(p_{a(ij1)} - p^{\text{sel}}_a - \Delta_{\text{repr}} p_a\right) \\ &\times \left(p_{a(ij2)} - p^{\text{sel}}_a - \Delta_{\text{repr}} p_a\right)\Big] \end{aligned} \tag{A24}$$

where $p^{\text{sel}}_m$, $p^{\text{sel}}_a$ are allele frequencies before reproduction (after selection). Expanding equation A24 yields:

$$\begin{aligned} D^{\text{repr}}_{ma,a} = {} & D^{\text{r}}_{ma,a} - \left(\Delta_{\text{repr}} p_m\right) D^{\text{r}}_{a,a} - \left(\Delta_{\text{repr}} p_a\right) D^{\text{r}}_{m,a} - \left(\Delta_{\text{repr}} p_a\right) D^{\text{r}}_{ma} \\ & + 2\left(\Delta_{\text{repr}} p_m\right)\left(\Delta_{\text{repr}} p_a\right)^2 \end{aligned} \tag{A25}$$

where associations $D^{\text{r}}_{ma,a}$, $D^{\text{r}}_{a,a}$, $D^{\text{r}}_{m,a}$ and $D^{\text{r}}_{ma}$ are measured among juveniles (after reproduction), but using alleles frequencies before reproduction ($p^{\text{sel}}_m$, $p^{\text{sel}}_a$) as "reference values" (Kirkpatrick et al., 2002), for example:

$$D^{\text{r}}_{ma,a} = \mathrm{E}^{\text{repr}}_{ij}\left[\left(p_{m(ij)} - p^{\text{sel}}_m\right)\left(p_{a(ij1)} - p^{\text{sel}}_a\right)\left(p_{a(ij2)} - p^{\text{sel}}_a\right)\right] \tag{A26}$$

$$D^{\mathrm{r}}_{a,a} = \mathrm{E}^{\mathrm{repr}}_{ij}\left[\left(p_{a(ij1)} - p^{\mathrm{sel}}_{a}\right)\left(p_{a(ij2)} - p^{\mathrm{sel}}_{a}\right)\right]. \tag{A27}$$

Because $\Delta_{\mathrm{repr}} p_m$ and $\Delta_{\mathrm{repr}} p_a$ are of order $\delta\sigma$, while associations $D^{\mathrm{r}}_{ma}$ and $D^{\mathrm{r}}_{m,a}$ are generated by the modifier effect and are thus also of order $\delta\sigma$, equation A25 simplifies to:

$$D^{\mathrm{repr}}_{ma,a} = D^{\mathrm{r}}_{ma,a} - \left(\Delta_{\mathrm{repr}} p_m\right) D^{\mathrm{r}}_{a,a} + o(\delta\sigma). \tag{A28}$$

Using the same reasoning as for $\Delta_{\mathrm{repr}} p_m$ above, $D^{\mathrm{r}}_{ma,a}$ is given by:

$$\begin{aligned} D^{\mathrm{r}}_{ma,a} = {} & \mathrm{E}^{\mathrm{sel}}_{ij}\left[\frac{1-\sigma_{ij}}{1-\sigma_i+\sigma_i/c}\,\zeta_{ma,a(ij)}\right] \\ & + \mathrm{E}^{\mathrm{sel}}_{ijk}\left[\frac{\sigma_{ij}\sigma_{ik}}{c\,\sigma_i\left[1-\sigma_i+\sigma_i/c\right]}\,\left[\left(1-r_{ma}\right)\zeta_{ma(ij)} + r_{ma}\,\zeta_{m,a(ij)}\right]\zeta_{a(ik)}\right]. \end{aligned} \tag{A29}$$

Using equations A16 – A17 and expressing the fractions in equation A29 to the first order in $\delta\sigma$ yields (after eliminating repeated indices from associations using equation A20):

$$\begin{aligned} D^{\mathrm{r}}_{ma,a} = {} & \left(1-\sigma_{\mathrm{e}}\right) D^{\mathrm{sel}}_{ma,a} + \sigma_{\mathrm{e}}\left(1-r_{ma}\right) D^{\mathrm{sel}}_{ma\overset{\frown}{/}a} + \sigma_{\mathrm{e}} r_{ma}\, D^{\mathrm{sel}}_{m,a\overset{\frown}{/}a} \\ & - c\left[\delta\sigma_{m,\mathrm{e}} + \left(1-2p_m\right)\delta\sigma_{m,m,\mathrm{e}}\right] D^{\mathrm{sel}}_{ma,ma} \\ & + \delta\sigma_{m,\mathrm{e}}\left[\left(1-r_{ma}\right) D^{\mathrm{sel}}_{ma\overset{\frown}{/}ma} + D^{\mathrm{sel}}_{ma\overset{\frown}{/}m,a} + r_{ma}\, D^{\mathrm{sel}}_{m,a\overset{\frown}{/}m,a}\right] \\ & - c\,\delta\sigma_{m,\mathrm{e}}\, p_m q_m\, D^{\mathrm{sel}}_{a,a} + \delta\sigma_{m,\mathrm{e}}\, p_m q_m\, D^{\mathrm{sel}}_{a\overset{\frown}{/}a} \end{aligned} \tag{A30}$$

with $\sigma_{\mathrm{e}} = \sigma/\left[c\left(1-\sigma\right)+\sigma\right]$. Several additional associations (such as $D_{ma,a\overset{\frown}{/}m}$, $D_{ma,m\overset{\frown}{/}a}$, $D_{ma\overset{\frown}{/}m\overset{\frown}{/}a}$) are generated from equation A29, but can be shown to be of order $1/N^2$. Finally, the term $-\left(\Delta_{\mathrm{repr}} p_m\right) D^{\mathrm{r}}_{a,a}$ in equation A28 can be computed as follows. Because $\Delta_{\mathrm{repr}} p_m$ is of order $\delta\sigma$, it is sufficient to evaluate $D^{\mathrm{r}}_{a,a}$ when $\delta\sigma = 0$, which is given by $\left(1-\sigma_{\mathrm{e}}\right) D^{\mathrm{sel}}_{a,a} + \sigma_{\mathrm{e}}\, D^{\mathrm{sel}}_{a\overset{\frown}{/}a}$. Furthermore, because $D^{\mathrm{r}}_{a,a}$ is of order $1/N$, it is sufficient to evaluate $\Delta_{\mathrm{repr}} p_m$ in the limit as $N$ tends to infinity, given by $-\left(c-1\right)\delta\sigma_{m,\mathrm{e}}\, p_m q_m$ (from equation A21). Therefore,

$$-\left(\Delta_{\mathrm{repr}} p_m\right) D^{\mathrm{r}}_{a,a} \approx \left(c-1\right)\delta\sigma_{m,\mathrm{e}}\, p_m q_m\left[\left(1-\sigma_{\mathrm{e}}\right) D^{\mathrm{sel}}_{a,a} + \sigma_{\mathrm{e}}\, D^{\mathrm{sel}}_{a\overset{\frown}{/}a}\right]. \tag{A31}$$

Equations A28, A30 and A31 yield:

$$\begin{aligned} D'_{ma,a} = D^{\mathrm{repr}}_{ma,a} = {} & \left(1-\sigma_{\mathrm{e}}\right) D^{\mathrm{sel}}_{ma,a} + \sigma_{\mathrm{e}}\left(1-r_{ma}\right) D^{\mathrm{sel}}_{ma\overset{\frown}{/}a} + \sigma_{\mathrm{e}} r_{ma}\, D^{\mathrm{sel}}_{m,a\overset{\frown}{/}a} \\ & - c\left[\delta\sigma_{m,\mathrm{e}} + \left(1-2p_m\right)\delta\sigma_{m,m,\mathrm{e}}\right] D^{\mathrm{sel}}_{ma,ma} \\ & + \delta\sigma_{m,\mathrm{e}}\left[\left(1-r_{ma}\right) D^{\mathrm{sel}}_{ma\overset{\frown}{/}ma} + D^{\mathrm{sel}}_{ma\overset{\frown}{/}m,a} + r_{ma}\, D^{\mathrm{sel}}_{m,a\overset{\frown}{/}m,a}\right] \\ & - \delta\sigma_{m,\mathrm{e}}\left[1+\left(c-1\right)\sigma_{\mathrm{e}}\right] p_m q_m\left(D^{\mathrm{sel}}_{a,a} - D^{\mathrm{sel}}_{a\overset{\frown}{/}a}\right). \end{aligned} \tag{A32}$$

Using the same reasoning as for the derivation of equation A25, we have:

$$D^{\mathrm{sel}}_{ma,a} = D^{\mathrm{s}}_{ma,a} - (\Delta_{\mathrm{sel}} p_m)\, D^{\mathrm{s}}_{a,a} - (\Delta_{\mathrm{sel}} p_a)\, D^{\mathrm{s}}_{m,a} - (\Delta_{\mathrm{sel}} p_a)\, D^{\mathrm{r}}_{ma} + 2\,(\Delta_{\mathrm{sel}} p_m)\,(\Delta_{\mathrm{sel}} p_a)^2 \tag{A33}$$

where $\Delta_{\mathrm{sel}} p_m$, $\Delta_{\mathrm{sel}} p_a$ are changes in allele frequencies during selection, and where $D^{\mathrm{s}}_{\mathbb{U},\mathbb{V}}$ are associations measured after selection (*i.e.*, weighting each parent by its relative fecundity), but using as reference values allele frequencies before selection. For example:

$$D^{\mathrm{s}}_{ma,a} = \mathrm{E}^{\mathrm{sel}}_{ij}\left[\left(p_{m(ij)} - p_m\right)\left(p_{a(ij1)} - p_a\right)\left(p_{a(ij2)} - p_a\right)\right]. \tag{A34}$$

Because the change in frequency of allele $m$ during selection ($\Delta_{\mathrm{sel}} p_m$) is generated by associations between the two loci (see equation A12), which are necessarily of order $p_a$ as they vanish when $p_a = 0$, both $\Delta_{\mathrm{sel}} p_m$ and $\Delta_{\mathrm{sel}} p_a$ are of order $p_a$, so that all terms on the right hand side of equation A33 except $D^{\mathrm{s}}_{ma,a}$ are of order ${p_a}^3$ or ${p_a}^2/N$, and can thus be neglected under our assumptions. We then have:

$$D^{\mathrm{s}}_{ma,a} = \mathrm{E}_{ij}\left[\frac{f_{ij}}{f_i}\,\zeta_{ma,a(ij)}\right]. \tag{A35}$$

Using equation A10, this gives:

$$D^{\mathrm{s}}_{ma,a} \approx -sh\left(D_{maa,a} + D_{ma,aa} - 2D_{ma,a\frown a}\right) - s\,(1-2h)\,(D_{maa,aa} - D_{ma,a\frown a,a}). \tag{A36}$$

Repeated indices can be eliminated using equation A20. For example, $D_{maa,a} = p_a q_a D_{m,a} + (1-2p_a)\, D_{ma,a}$; however, because $D_{m,a}$ and $D_{ma,a}$ are both of order $p_a$, this is equivalent to $D_{ma,a}$ to leading order in $p_a$. Similarly, $D_{ma,aa}$ and $D_{maa,aa}$ are both equivalent to $D_{ma,a}$, while $D_{ma,a\frown a}$ and $D_{ma,a\frown a,a}$ can be shown to be of order $1/N^2$. Therefore,

$$D^{\mathrm{sel}}_{ma,a} \approx (1-s)\, D_{ma,a}\,. \tag{A37}$$

Taking into account the term shown in equation A11 introduces associations that are of order $1/N^2$, so that equation A37 should hold for arbitrary $s$, provided that $p_a$ remains small. The effect of selection on the other associations appearing in equation A32 is obtained similarly. In particular, we have:

$$D^{\mathrm{sel}}_{ma\frown a} \approx D^{\mathrm{s}}_{ma\frown a} = \mathrm{E}_{ijk}\left[\left(\frac{f_{ij}}{f_i}\,\zeta_{ma(ij)}\right)\left(\frac{f_{ik}}{f_i}\,\zeta_{a(ik)}\right)\right] \approx (1-2sh)\, D_{ma\frown a}, \tag{A38}$$

$$D^{\mathrm{sel}}_{ma\frown a} \approx (1-2sh)\, D_{ma\frown a}, \qquad D^{\mathrm{sel}}_{ma\frown ma} \approx (1-2sh)\, D_{ma\frown ma}, \tag{A39}$$

$$D^{\text{sel}}_{ma\overset{\frown}{/}m,a} \approx (1-2sh)\, D_{ma\overset{\frown}{/}m,a}, \qquad D^{\text{sel}}_{m,a\overset{\frown}{/}m,a} \approx (1-2sh)\, D_{m,a\overset{\frown}{/}m,a}, \tag{A40}$$

$$D^{\text{sel}}_{ma,ma} \approx (1-s)\, D_{ma,ma}\,. \tag{A41}$$

Expressions for $D^{\text{sel}}_{a,a}$ and $D^{\text{sel}}_{a\overset{\frown}{/}a}$ (that also appear in equation A32) are derived in the next subsection.

**Recursions on $\boldsymbol{D_{a,a}}$ and $\boldsymbol{D_{a\overset{\frown}{/}a}}$.** As shown by equation A32, the association $D_{ma,a}$ (involved in the change in frequency of the modifier) is generated in part by associations $D_{a,a}$ and $D_{a\overset{\frown}{/}a}$. However, because these associations are multiplied by $\delta\sigma_{m,\text{e}}$ in equation A32, it is sufficient to express them when $\delta\sigma = 0$. The association $D_{a,a}$ remains unchanged during dispersal and drift within demes: $D'_{a,a} = D^{\text{disp}}_{a,a} = D^{\text{repr}}_{a,a}$. Neglecting the modifier effect, we have:

$$D^{\text{repr}}_{a,a} = (1-\sigma_{\text{e}})\, D^{\text{sel}}_{a,a} + \sigma_{\text{e}}\, D^{\text{sel}}_{a\overset{\frown}{/}a}\,. \tag{A42}$$

We then have:

$$D^{\text{sel}}_{a,a} = D^{\text{s}}_{a,a} - (\Delta_{\text{sel}} p_a)^2\,, \qquad D^{\text{sel}}_{a\overset{\frown}{/}a} = D^{\text{s}}_{a\overset{\frown}{/}a} - (\Delta_{\text{sel}} p_a)^2 \tag{A43}$$

with

$$D^{\text{s}}_{a,a} = \text{E}_{ij}\left[\frac{f_{ij}}{f_i}\,\zeta_{a,a(ij)}\right], \quad D^{\text{s}}_{a\overset{\frown}{/}a} = \text{E}_{ijk}\left[\left(\frac{f_{ij}}{f_i}\,\zeta_{a(ij)}\right)\left(\frac{f_{ik}}{f_i}\,\zeta_{a(ik)}\right)\right]. \tag{A44}$$

Using equations A10 and A20, and given that associations $D_{a,a\overset{\frown}{/}a}$ and $D_{a,a\overset{\frown}{/}a,a}$ are of order $1/N^2$, one obtains, to the first order in $s$:

$$\begin{aligned} D^{\text{s}}_{a,a} &\approx (1-s)\, D_{a,a} - s\,(1-2h)\,(p_a q_a)^2 \\ &\approx (1-s)\, D_{a,a} - s\,(1-2h)\, {p_a}^2 \end{aligned} \tag{A45}$$

$$D^{\text{s}}_{a\overset{\frown}{/}a} \approx (1-2sh)\, D_{a\overset{\frown}{/}a}\,. \tag{A46}$$

For arbitrary $s$ (but to leading order in $p_a$), equation A45 stays unchanged, as the term given by equation A11 generates associations that are of order $1/N^2$; however, equation A46 becomes:

$$D^{\text{s}}_{a\overset{\frown}{/}a} \approx (1-2sh)\, D_{a\overset{\frown}{/}a} + (shp_a)^2\,. \tag{A47}$$

Finally, since $\Delta_{\mathrm{sel}} p_a$ is of order $p_a$, it is sufficient to express the term $(\Delta_{\mathrm{sel}} p_a)^2$ in equation A43 in the limit as deme size tends to infinity (as taking finite deme size into account would introduce terms of order ${p_a}^2/N$), that is, $(\Delta_{\mathrm{sel}} p_a)^2 \approx (shp_a)^2$. This finally gives:

$$D^{\mathrm{sel}}_{a,a} \approx (1-s)\, D_{a,a} - s\left(1 - 2h + h^2 s\right) {p_a}^2 \tag{A48}$$

$$D^{\mathrm{sel}}_{\widehat{a/a}} \approx (1-2sh)\, D_{\widehat{a/a}}\,. \tag{A49}$$

Contrarily to $D_{a,a}$, the association $D_{\widehat{a/a}}$ is affected by dispersal and drift within demes. The effect of drift can be obtained by expressing associations involving individuals sampled with replacement from a deme in terms of associations involving individuals sampled without replacement. In particular, from equation A5 we have:

$$D_{\widehat{a/a}} = \frac{1}{2N}\left(p_a q_a + D_{a,a}\right) + \left(1 - \frac{1}{N}\right) D_{a/a}\,. \tag{A50}$$

However, the terms $D_{a,a}/N$ and $D_{a/a}/N$ are of order ${p_a}^2/N$ or $1/N^2$, so that:

$$D_{\widehat{a/a}} \approx \frac{p_a}{2N} + D_{a/a}\,. \tag{A51}$$

$D_{a/a}$ is the association between two $a$ alleles sampled from two different adults from the same deme; these alleles were necessarily in two different juveniles from the same deme just before drift, so that this association is the same as $D^{\mathrm{disp}}_{a/a}$ (measured among juveniles, just after dispersal). The effect of drift within demes is thus given by:

$$D_{\widehat{a/a}}{}' \approx \frac{p_a}{2N} + D^{\mathrm{disp}}_{a/a}\,. \tag{A52}$$

The effect of dispersal on $D_{a/a}$ is obtained as follows. With probability $(1-d)^2$, the two juveniles were already in the same deme before dispersal, so that the association becomes $D^{\mathrm{repr}}_{a/a}$. With probability $1-(1-d)^2$, the two juveniles come from different demes. In this case, their genetic compositions become independent (since the number of demes is infinite), so that the association becomes $(D^{\mathrm{repr}}_{a})^2 = 0$. Therefore,

$$D^{\mathrm{disp}}_{a/a} = (1-d)^2\, D^{\mathrm{repr}}_{a/a}\,. \tag{A53}$$

Finally, the association between two alleles in two juveniles is the same as the association among those alleles in two parents sampled with replacement from the same deme (after selection), so that:

$$D^{\mathrm{repr}}_{a/a} = D^{\mathrm{sel}}_{\widehat{a/a}} = (1-2sh)\, D_{\widehat{a/a}}\,. \tag{A54}$$

Altogether, recursions for $D_{a,a}$ and $D_{\widehat{a/a}}$ are thus given by:

$$D_{a,a}' \approx (1-\sigma_{\mathrm{e}})\left[(1-s)\,D_{a,a} + \iota\, {p_a}^2\right] + \sigma_{\mathrm{e}}\,(1-2sh)\,D_{\widehat{a/a}} \tag{A55}$$

$$D_{\widehat{a/a}}' \approx \frac{p_a}{2N} + (1-d)^2\,(1-2sh)\,D_{\widehat{a/a}} \tag{A56}$$

with $\iota = -s\,(1-2h+h^2 s)$. Equations A55 – A56 show that the association $D_{a,a}$ (measuring departure from Hardy-Weinberg equilibrium at the whole metapopulation level) is generated by two effects: an effect of selection in the presence of dominance at the selected locus, measured as a deviation from multiplicativity ($\iota \neq 0$, Otto, 2003), and an effect of population structure, through a genetic correlation between uniting gametes (term in $D_{\widehat{a/a}}$).

**Quasi-equilibrium approximation.** Once the frequency of the deleterious allele $a$ in the metapopulation has reached mutation-selection balance, all changes in allele frequencies are solely due to the modifier effect $\delta\sigma$, and are thus slow when $\delta\sigma$ is small. Because genetic associations are broken by dispersal, sex and recombination (as can be seen from equations A32, A55, A56), a separation of timescales argument can be used when $\delta\sigma \ll d$, $\sigma_{\mathrm{e}}$, $r_{ma}$, as genetic associations will reach a quasi-equilibrium value on a relatively short timescale compared to the change in allele frequencies (Nagylaki, 1993; Barton and Turelli, 1991; Roze and Rousset, 2008). In this case, associations can be expressed in terms of allele frequencies, by setting $D_{\mathcal{S}}' = D_{\mathcal{S}}$ and solving for $D_{\mathcal{S}}$. In the absence of modifier effect ($\delta\sigma = 0$), allele frequencies and genetic associations reach a proper equilibrium; in particular, equations A55 and A56 yield for $D_{a,a}$ and $D_{\widehat{a/a}}$ at equilibrium:

$$D_{\widehat{a/a}} \approx \frac{p_a}{2N\left[1-(1-d)^2\,(1-2sh)\right]} \tag{A57}$$

$$D_{a,a} \approx \frac{(1-\sigma_{\mathrm{e}})\,\iota\,{p_a}^2 + \sigma_{\mathrm{e}}\,(1-2sh)\,D_{\widehat{a/a}}}{1-(1-\sigma_{\mathrm{e}})\,(1-s)}\,. \tag{A58}$$

Equation A58 is equivalent to equation 8 in Otto (2003) when deme size $N$ tends to infinity.
