## Supplementary material for "Deleterious mutations and selection for sex in spatially structured, diploid populations": File S2

FILE S2: RECURSIONS IN THE TWO-LOCUS MODEL

We provide here recurrence equations for genetic associations involved in the change in frequency of the modifier in the two-locus model, as well as some interpretations for these associations and the mechanisms that generate them. The recursions can be derived using the methods explained in File S1, that have been implemented in the *Mathematica* notebook available as Supplementary Material.

**Associations generated by population structure and selection.** Indirect selection on the sex modifier stems from its effect on genetic associations that are themselves generated by population structure and selection against the deleterious allele. In order to obtain an approximation for indirect selection to the first order in $\delta\sigma$, it is sufficient to express these associations when $\delta\sigma = 0$ (*i.e.*, neglecting the effect of the modifier).

*Associations* $D_{a,a}$ *and* $D_{a\widehat{/}a}$. At the selected locus, population structure and selection generate the pairwise associations $D_{a,a}$ and $D_{a\widehat{/}a}$. As shown in File S1, recursions for these associations are given by:

$$D_{a,a}' \approx (1-\sigma_{\mathrm{e}}) \left[(1-s)\, D_{a,a} + \iota\, {p_a}^2\right] + \sigma_{\mathrm{e}}\, (1-2sh)\, D_{a\widehat{/}a} \tag{B1}$$

$$D_{a\widehat{/}a}' \approx \frac{p_a}{2N} + (1-d)^2\, (1-2sh)\, D_{a\widehat{/}a} \tag{B2}$$

with $\iota = -s\,(1-2h+h^2 s)$ and $\sigma_{\mathrm{e}} = \sigma / \left[c\,(1-\sigma)+\sigma\right]$, giving at equilibrium:

$$D_{a\widehat{/}a} \approx \frac{p_a}{2N\left[1-(1-d)^2\,(1-2sh)\right]} \tag{B3}$$

$$D_{a,a} \approx \frac{(1-\sigma_{\mathrm{e}})\,\iota\,{p_a}^2 + \sigma_{\mathrm{e}}\,(1-2sh)\, D_{a\widehat{/}a}}{1-(1-\sigma_{\mathrm{e}})\,(1-s)}\,. \tag{B4}$$

This yields:

$$\begin{aligned} D_{a,a} - D_{a\widehat{/}a} \approx\; & \frac{(1-\sigma_{\mathrm{e}})\,\iota\,{p_a}^2}{1-(1-\sigma_{\mathrm{e}})\,(1-s)} \\ & - \frac{s\left[1-(1-2h)\,\sigma_{\mathrm{e}}\right] p_a}{2N\left[1-(1-d)^2\,(1-2sh)\right]\left[1-(1-\sigma_{\mathrm{e}})\,(1-s)\right]}\,. \end{aligned} \tag{B5}$$

When $d$ and $s$ are small, this is approximately:

$$D_{a\widehat{/}a} \approx \frac{p_a}{4N\,(d+sh)}, \qquad D_{a,a} \approx \frac{(1-\sigma_{\mathrm{e}})\,\iota\,{p_a}^2}{\sigma_{\mathrm{e}}+s} + \frac{\sigma_{\mathrm{e}}\,p_a}{4N\,(d+sh)\,(\sigma_{\mathrm{e}}+s)}, \tag{B6}$$

$$D_{a,a} - D_{\widehat{a/a}} \approx \frac{(1-\sigma_{\mathrm{e}})\,\iota\, {p_a}^2}{\sigma_{\mathrm{e}} + s} - \frac{s\,[1-(1-2h)\,\sigma_{\mathrm{e}}]\,p_a}{4N\,(d+sh)\,(\sigma_{\mathrm{e}}+s)}\,. \tag{B7}$$

$D_{\widehat{a/a}}$ and $D_{a,a}$ can be shown to be equivalent to $F_{\mathrm{ST}}\, p_a$ and $F_{\mathrm{IT}}\, p_a$, respectively ($p_a$ being replaced by $p_a q_a$ in these expressions when $p_a$ is not small; e.g., Roze and Rousset, 2008; Roze, 2015). Note that the approximation given by equation B6 for $F_{\mathrm{ST}}$ differs slightly from the expression $1/\left[1+4N\,(d+sh)\right]$ obtained by Glémin et al. (2003) and Roze (2015); this is due to the fact that our derivations assume that $1/N$ is much smaller than $d$, $s$ and $\sigma_{\mathrm{e}}$ (so that $Nd$, $Ns \gg 1$). Finally, because $F_{\mathrm{IS}} = (F_{\mathrm{IT}} - F_{\mathrm{ST}})\,/\,(1-F_{\mathrm{ST}})$, the difference $D_{a,a} - D_{\widehat{a/a}}$ provides an approximation for $F_{\mathrm{IS}}\, p_a$ (neglecting terms in ${p_a}^2/N$ and in $1/N^2$). Equations B5 and B7 show that $F_{\mathrm{IS}}$ is generated by a departure from multiplicative fitness effects at the selected locus, provided that reproduction is partially clonal (term in $\iota$), and by an effect of selection and population structure that tends to generate negative $F_{\mathrm{IS}}$ (term in $s/N$). This last effect can be understood from equation B1. Neglecting terms in ${p_a}^2$, $D_{a,a}$ corresponds approximately to the frequency of $aa$ homozygotes in the metapopulation, while $D_{\widehat{a/a}}$ is approximately the probability of drawing two $a$ alleles by sampling two genes with replacement from the same deme, both quantities being measured among newly formed adults (before selection). In the absence of selection, these quantities are equal and are not affected by the rate of clonal reproduction. However, selection tends to reduce $D_{a,a}$ relative to $D_{\widehat{a/a}}$ through two different effects: (i) when reproduction is partially clonal, selection tends to reduce the frequency of $aa$ individuals produced clonally (term in $(1-\sigma_{\mathrm{e}})\,(1-s)$ in equation B1); (ii) selection against allele $a$ reduces the probability that two uniting gametes both carry $a$, compared to the probability of drawing two $a$ alleles among adults before selection (term in $\sigma_{\mathrm{e}}\,(1-2sh)$ in equation B1). As a result, the frequency of $aa$ homozygotes among newly formed adults tends to be lower than the probability of drawing two $a$ alleles when sampling with replacement from those adults.

*Associations $D_{ma,ma}$, $D_{\widehat{ma/ma}}$, $D_{\widehat{ma/m,a}}$ and $D_{\widehat{m,a/m,a}}$.* These associations are generated by population structure. Recursions are given by:

$$D_{\widehat{m,a/m,a}}{}' \approx \frac{p_m q_m p_a}{2N} + (1-d)^2\,(1-\sigma_{\mathrm{e}})^2\,(1-2sh)\,D_{\widehat{m,a/m,a}} \tag{B8}$$

$$D_{ma\overset{\frown}{/}m,a}{}' \approx (1-d)^2 (1-\sigma_{\rm e}) (1-2sh) \left[(1-\sigma_{\rm e} r_{ma}) D_{ma\overset{\frown}{/}m,a} + \sigma_{\rm e} r_{ma} D_{m,a\overset{\frown}{/}m,a}\right] \quad \text{(B9)}$$

$$\begin{aligned} D_{ma\overset{\frown}{/}ma}{}' \approx \frac{p_m q_m p_a}{2N} &+ (1-d)^2 (1-2sh) \Big[(1-\sigma_{\rm e} r_{ma})^2 D_{ma\overset{\frown}{/}ma} \\ &+ 2\sigma_{\rm e} r_{ma} (1-\sigma_{\rm e} r_{ma})^2 D_{ma\overset{\frown}{/}m,a} + (\sigma_{\rm e} r_{ma})^2 D_{m,a\overset{\frown}{/}m,a}\Big] \end{aligned} \quad \text{(B10)}$$

$$\begin{aligned} D_{ma,ma}{}' \approx (1-\sigma_{\rm e}) (1-s) D_{ma,ma} &+ \sigma_{\rm e} (1-2sh) \Big[(1-r_{ma})^2 D_{ma\overset{\frown}{/}ma} \\ &+ 2r_{ma} (1-r_{ma})^2 D_{ma\overset{\frown}{/}m,a} + r_{ma}{}^2 D_{m,a\overset{\frown}{/}m,a}\Big] \end{aligned} \quad \text{(B11)}$$

Solving equations B8 – B11 for $D_{\mathcal{S}}{}' = D_{\mathcal{S}}$ yields expressions for these different associations at quasi-equilibrium. When $s$, $d$ and $\sigma_{\rm e}$ are small, these are approximately:

$$D_{m,a\overset{\frown}{/}m,a} \approx \frac{p_m q_m p_a}{4N (d + sh + \sigma_{\rm e})} \quad \text{(B12)}$$

$$D_{ma\overset{\frown}{/}m,a} \approx \frac{\sigma_{\rm e} r_{ma} p_m q_m p_a}{4N (d + sh + \sigma_{\rm e}) (2d + 2sh + \sigma_{\rm e} + \sigma_{\rm e} r_{ma})} \quad \text{(B13)}$$

$$D_{ma\overset{\frown}{/}ma} \approx \frac{p_m q_m p_a}{4N (d + sh + \sigma_{\rm e} r_{ma})} \left[1 + \frac{(\sigma_{\rm e} r_{ma})^2}{(d + sh + \sigma_{\rm e}) (2d + 2sh + \sigma_{\rm e} + \sigma_{\rm e} r_{ma})}\right] \quad \text{(B14)}$$

$$D_{ma,ma} \approx \frac{[1 - 2r_{ma} (1 - r_{ma})] \sigma_{\rm e} p_m q_m p_a}{4N (s + \sigma_{\rm e}) (d + sh + \sigma_{\rm e} r_{ma})} \,. \quad \text{(B15)}$$

The association $D_{ma\overset{\frown}{/}ma}$ is related to the variance in linkage disequilibrium (LD) among demes. Indeed, the LD within deme $i$ can be defined as $D_{ma(i)} = p_{ma(i)} - p_{m(i)} p_{a(i)}$ where $p_{ma(i)}$ is the frequency of $ma$ haplotypes in deme $i$. When $\delta\sigma = 0$, the average LD should be zero at equilibrium, so that the variance in LD among demes is $\mathrm{E}_i \left[\left(p_{ma(i)} - p_{m(i)} p_{a(i)}\right)^2\right]$, which can be shown to be equivalent to $D_{ma\overset{\frown}{/}ma} - 2D_{ma\overset{\frown}{/}m\overset{\frown}{/}a} + D_{m\overset{\frown}{/}m\overset{\frown}{/}a\overset{\frown}{/}a}$. However, associations $D_{ma\overset{\frown}{/}m\overset{\frown}{/}a}$ and $D_{m\overset{\frown}{/}m\overset{\frown}{/}a\overset{\frown}{/}a}$ are of order $1/N^2$ at equilibrium, so that the variance in LD among demes is approximately equal to $D_{ma\overset{\frown}{/}ma}$. As shown by equation B10, it is generated by drift within demes (term in $1/N$), which produces either positive or negative LD within each deme. Similarly, drift generates a variance among demes in the association between $m$ and $a$ on different haplotypes of the same individual ($D_{m,a(i)}$), measured by $D_{m,a\overset{\frown}{/}m,a}$ (equation B8). Recombination converts this variance in $D_{m,a(i)}$ into a covariance between $D_{m,a(i)}$ and $D_{ma(i)}$ and into a variance in $D_{ma(i)}$, as shown by equations B9 and B10.

Finally, the association $D_{ma,ma}$ is positive when homozygotes at the modifier locus tend to be also homozygous at the selected locus, and negative when homozygotes at the modifier locus tend to be heterozygous at the selected locus (indeed, genotypes that are homozygous at both loci or heterozygous at both loci contribute positively to $D_{ma,ma}$, while genotypes that are homozygous at one locus and heterozygous at the other contribute negatively). As shown by equation B11, this association is positive and is generated by the variance between demes in LD among gametes. For example, under extreme negative or positive LD (*i.e.*, when only $mA$ or $Ma$ gametes are present, or when only $ma$ or $MA$ gametes are present), only double homozygotes or double heterozygotes are produced (while $D_{ma,ma}$ among offspring is zero in the absence of LD and when mating is random).

*Associations $D_{ma,m}$, $D_{ma\widehat{/}m}$ and $D_{m,a\widehat{/}m}$.* These associations are generated by population structure and selection against allele $a$. Recursions are given by:

$$D_{m,a\widehat{/}m}{}' \approx (1-d)^2\,(1-\sigma_{\mathrm{e}})\left[(1-sh)\,D_{m,a\widehat{/}m} - sh\left(D_{ma\widehat{/}m,a} + D_{m,a\widehat{/}m,a}\right)\right] \tag{B16}$$

$$\begin{aligned} D_{ma\widehat{/}m}{}' \approx (1-d)^2 \Big[&(1-sh)\left[(1-\sigma_{\mathrm{e}} r_{ma})\,D_{ma\widehat{/}m} + \sigma_{\mathrm{e}} r_{ma} D_{m,a\widehat{/}m}\right] \\ &- sh\left[(1-\sigma_{\mathrm{e}} r_{ma})\,D_{ma\widehat{/}ma} + D_{ma\widehat{/}m,a} + \sigma_{\mathrm{e}} r_{ma} D_{m,a\widehat{/}m,a}\right]\Big] \end{aligned} \tag{B17}$$

$$\begin{aligned} D_{ma,m}{}' \approx\ &(1-\sigma_{\mathrm{e}})\left[(1-sh)\,D_{ma,m} - s\,(1-h)\,D_{ma,ma}\right] \\ &+ \sigma_{\mathrm{e}}\,(1-sh)\left[(1-r_{ma})\,D_{ma\widehat{/}m} + r_{ma} D_{m,a\widehat{/}m}\right] \\ &- \sigma_{\mathrm{e}} sh\left[(1-r_{ma})\,D_{ma\widehat{/}ma} + D_{ma\widehat{/}m,a} + r_{ma} D_{m,a\widehat{/}m,a}\right]. \end{aligned} \tag{B18}$$

When $s$, $d$, $\sigma_{\mathrm{e}}$ and $r_{ma}$ are small, the quasi-equilibrium solutions are approximately:

$$D_{m,a\widehat{/}m} \approx -\frac{sh\,(2d+2sh+\sigma_{\mathrm{e}})\,p_m q_m p_a}{4N\,(d+sh+\sigma_{\mathrm{e}})\,(2d+sh+\sigma_{\mathrm{e}})\,Z} \tag{B19}$$

$$D_{ma\widehat{/}m} \approx -\frac{sh\,(2d+2sh+\sigma_{\mathrm{e}})\,p_m q_m p_a}{4N\,(d+sh+\sigma_{\mathrm{e}} r_{ma})\,(2d+sh+\sigma_{\mathrm{e}} r_{ma})\,Z} \tag{B20}$$

$$D_{ma,m} \approx -\frac{\left[s\,(1-h)\,(2d+sh) + sh\,(s+\sigma_{\mathrm{e}})\right]\sigma_{\mathrm{e}}\,(2d+2sh+\sigma_{\mathrm{e}})\,p_m q_m p_a}{4N\,(s+\sigma_{\mathrm{e}})\,(sh+\sigma_{\mathrm{e}})\,(d+sh+\sigma_{\mathrm{e}} r_{ma})\,(2d+sh+\sigma_{\mathrm{e}} r_{ma})\,Z} \tag{B21}$$

with $Z = 2d + 2sh + \sigma_{\mathrm{e}} + \sigma_{\mathrm{e}} r_{ma}$ (note that the $\sigma_{\mathrm{e}} r_{ma}$ products in the denominators should be negligible when $r_{ma}$ and $\sigma_{\mathrm{e}}$ are both small; however, keeping these terms improves the approximations when $r_{ma}$ is not very small). More accurate expressions for arbitrary $\sigma_{\mathrm{e}}$ and $r_{ma}$ can be obtained from equations B16 – B18.

The association $D_{ma\widehat{/}m}$ is approximately equal to the covariance between the within-deme linkage disequilibrium $D_{ma(i)}$ (as defined above) and the frequency of allele

$m$ in deme $i$, $p_{m(i)}$. Indeed, given that the average LD should be zero when $\delta\sigma = 0$, this covariance can be shown to be equal to $D_{ma\overset{\frown}{/}m} - D_{m\overset{\frown}{/}m\overset{\frown}{/}a}$, while $D_{m\overset{\frown}{/}m\overset{\frown}{/}a}$ is of order $1/N^2$ at equilibrium. Similarly, $D_{m,a\overset{\frown}{/}m}$ is approximately equal to the covariance between $D_{m,a(i)}$ and $p_{m(i)}$. As shown the equations B16 and B17, these covariances are negative and are generated by the variances in $D_{ma(i)}$ and $D_{m,a(i)}$ among demes (represented by associations $D_{ma\overset{\frown}{/}ma}$ and $D_{m,a\overset{\frown}{/}m,a}$, as explained above). Indeed, when $D_{ma(i)}$ and/or $D_{m,a(i)}$ happen to be positive in a given deme (due to drift), allele $m$ is associated with the deleterious allele $a$, causing a decrease in frequency of allele $m$ in this deme (conversely, the frequency of $m$ within the deme tends to increase when $D_{ma(i)}$ and/or $D_{m,a(i)}$ are negative).

Finally, the association $D_{ma,m}$ is negative when allele $a$ tends to be more frequent among heterozygotes at the modifier locus than among homozygotes, and positive when $a$ is more frequent among homozygotes. As shown by equation B18, two different effects generate negative $D_{ma,m}$. The first effect involves the association $D_{ma,ma}$, representing the correlation in homozygosity between the two loci: indeed, because homozygotes at the modifier locus tend to be also homozygous at the selected locus (see above), selection against allele $a$ is more efficient among homozygotes at the modifier locus than among heterozygotes, causing a higher frequency of $a$ among heterozygotes at the modifier locus than among homozygotes. The second effect is generated by the negative covariances between $D_{ma(i)}$ and $p_{m(i)}$, and between $D_{m,a(i)}$ and $p_{m(i)}$. As discussed in the previous paragraph, these covariances are negative; therefore, when allele $a$ tends to be associated with allele $m$ within a deme ($D_{ma(i)} > 0$), the frequency of $m$ in the deme tends to be relatively lower, so that $ma$ gametes have a relatively higher chance of fusing with a gamete carrying allele $M$, generating an association between allele $a$ and heterozygosity at the modifier locus. Conversely, when $a$ tends to be associated with $M$ ($D_{ma(i)} < 0$), the frequency of $m$ is relatively higher, so that $Ma$ gametes have an increased probability of fusing with an $m$ gamete (also generating an association between allele $a$ and heterozygosity at the modifier locus).

**Associations generated by the modifier effect.** Indirect selection at the modifier locus involves associations $D_{ma}$, $D_{m,a}$, $D_{m\widehat{/}a}$ and $D_{ma,a}$ (see equation A12 in File S1), that are generated by the effect of the sex modifier on the associations described above.

*Associations $D_{ma,a}$, $D_{ma\widehat{/}a}$, $D_{m,a\widehat{/}a}$ and $D_{m\widehat{/}a,a}$.* The association $D_{m\widehat{/}a,a}$ is found to be zero to the first order in $1/N$, while recursions on the other associations are given by:

$$D_{m,a\widehat{/}a}{}' \approx (1-d)^2 (1-\sigma_{\mathrm{e}}) (1-2sh) \left[D_{m,a\widehat{/}a} - d\sigma_{m,\mathrm{e}} (c-1) \left(D_{ma\widehat{/}m,a} + D_{m,a\widehat{/}m,a}\right)\right] \tag{B22}$$

$$\begin{aligned} D_{ma\widehat{/}a}{}' \approx (1-d)^2 (1-2sh) &\Big[(1-\sigma_{\mathrm{e}} r_{ma}) D_{ma\widehat{/}a} + \sigma_{\mathrm{e}} r_{ma} D_{m,a\widehat{/}a} \\ &- d\sigma_{m,\mathrm{e}} (c-1) \left[(1-\sigma_{\mathrm{e}} r_{ma}) D_{ma\widehat{/}ma} + D_{ma\widehat{/}m,a} + \sigma_{\mathrm{e}} r_{ma} D_{m,a\widehat{/}m,a}\right]\Big] \end{aligned} \tag{B23}$$

$$\begin{aligned} D_{ma,a}{}' \approx (1-s) &\left[(1-\sigma_{\mathrm{e}}) D_{ma,a} - c\left[d\sigma_{m,\mathrm{e}} + d\sigma_{m,m,\mathrm{e}} (1-2p_m)\right] D_{ma,ma}\right] \\ &- d\sigma_{m,\mathrm{e}} \left[1 + (c-1)\sigma_{\mathrm{e}}\right] \left[(1-s) D_{a,a} - (1-2sh) D_{a\widehat{/}a}\right] p_m q_m \\ &+ (1-2sh) \Big[\sigma_{\mathrm{e}} (1-r_{ma}) D_{ma\widehat{/}a} + \sigma_{\mathrm{e}} r_{ma} D_{m,a\widehat{/}a} \\ &\quad + d\sigma_{m,\mathrm{e}} \left[(1-r_{ma}) D_{ma\widehat{/}ma} + D_{ma\widehat{/}m,a} + r_{ma} D_{m,a\widehat{/}m,a}\right]\Big] \end{aligned} \tag{B24}$$

with:

$$d\sigma_{m,\mathrm{e}} = \frac{\delta\sigma \left[h_m + (1-2h_m) p_m\right]}{c(1-\sigma)+\sigma}, \quad d\sigma_{m,m,\mathrm{e}} = \frac{\delta\sigma (1-2h_m)}{c(1-\sigma)+\sigma}. \tag{B25}$$

When $s$, $d$, $\sigma_{\mathrm{e}}$ and $r_{ma}$ are small, and $h_m = 1/2$ (additive modifier), the quasi-equilibrium solutions are approximately:

$$D_{m,a\widehat{/}a} \approx -\frac{d\sigma_{m,\mathrm{e}} (c-1) p_m q_m p_a}{4N (d+sh+\sigma_{\mathrm{e}}) (2d+2sh+\sigma_{\mathrm{e}}+\sigma_{\mathrm{e}} r_{ma})} \tag{B26}$$

$$D_{ma\widehat{/}a} \approx -\frac{d\sigma_{m,\mathrm{e}} (c-1) (2d+2sh+\sigma_{\mathrm{e}}) p_m q_m p_a}{4N (d+sh+\sigma_{\mathrm{e}} r_{ma}) (2d+2sh+\sigma_{\mathrm{e}} r_{ma}) (2d+2sh+\sigma_{\mathrm{e}}+\sigma_{\mathrm{e}} r_{ma})} \tag{B27}$$

$$\begin{aligned} D_{ma,a} \approx\ & \frac{d\sigma_{m,\mathrm{e}} (2d+2sh+\sigma_{\mathrm{e}}) \left[4s(d+sh) - (c-1)\sigma_{\mathrm{e}} (\sigma_{\mathrm{e}}+2d+s+2sh)\right] p_m q_m p_a}{4N (s+\sigma_{\mathrm{e}})^2 (d+sh+\sigma_{\mathrm{e}} r_{ma}) (2d+2sh+\sigma_{\mathrm{e}} r_{ma}) (2d+2sh+\sigma_{\mathrm{e}}+\sigma_{\mathrm{e}} r_{ma})} \\ & - \frac{d\sigma_{m,\mathrm{e}} \left[1+(c-1)\sigma_{\mathrm{e}}\right] \iota\, p_m q_m p_a{}^2}{(s+\sigma_{\mathrm{e}})^2}. \end{aligned} \tag{B28}$$

In the presence of a direct cost of sex ($c > 1$) and when allele $m$ increases investment in sex ($d\sigma_{m,\mathrm{e}} > 0$), the same mechanism that was generating negative values of $D_{m,a\widehat{/}m}$, $D_{ma\widehat{/}m}$ and $D_{ma,m}$ (and that was resulting from population structure and from the effects of selection against allele $a$, see equations B16 – B21 above) also tends to generate negative values of $D_{m,\widehat{a/a}}$, $D_{\widehat{ma/a}}$ and $D_{ma,a}$, since allele $m$ is deleterious (terms in $c-1$ in equations B22 – B24). As shown by equation B24, other sources of $D_{ma,a}$ stem from the effect of the sex modifier, and persist even when $c=1$ (no cost of sex). In particular, increasing sex tends to break the excess heterozygosity within demes at the selected locus ($D_{a,a} - D_{\widehat{a/a}} < 0$) by increasing the frequency of homozygotes, causing an association between the modifier allele increasing sex ($m$ if $d\sigma_{m,\mathrm{e}} > 0$) and homozygosity at the selected locus ($D_{ma,a} > 0$, second line of equation B24). The variance in linkage disequilibrium within demes also tends to generate positive $D_{ma,a}$ when $d\sigma_{m,\mathrm{e}} > 0$ (last line of equation B24). Indeed, uniting gametes have a relatively higher chance to carry the modifier allele increasing sex, in which case they also tend to carry the allele at the selected locus that is locally associated with this modifier allele, generating new homozygotes at the selected locus. Conversely, the correlation in homozygosity among loci tends to generate a negative association between the modifier allele increasing sex and homozygosity at the selected locus (term in $D_{ma,ma}$ on the first line of equation B24). Indeed, when $d\sigma_{m,\mathrm{e}} > 0$, sexual reproduction between $mmaa$ and $mmAA$ individuals generates an association between $m$ and $Aa$ (while $MMaa$ and $MMAA$ individuals engage less frequently in sex). When $c=1$ (no cost of sex), one can show that $D_{ma,a}$ is positive when allele $m$ increases the rate of sex; however, the sign of $D_{ma,a}$ may change when $c>1$ (due to the effect of direct selection against allele $m$), in particular when the rate of sex is large.

*Associations $D_{ma}$, $D_{m,a}$ and $D_{\widehat{m/a}}$.* Recursions on these associations are given by:

$$D_{\widehat{m/a}}{}' \approx (1-d)^2 \left[D^{\mathrm{sel}}_{\widehat{m/a}} - d\sigma_{m,\mathrm{e}}\,(c-1)\left(D^{\mathrm{sel}}_{\widehat{ma/m}} + D^{\mathrm{sel}}_{m,\widehat{a/m}}\right)\right] \tag{B29}$$

$$\begin{aligned} D_{m,a}{}' \approx\; & (1-\sigma_{\mathrm{e}})\,D^{\mathrm{sel}}_{m,a} + \sigma_{\mathrm{e}}\,D^{\mathrm{sel}}_{\widehat{m/a}} + d\sigma_{m,\mathrm{e}}\,D^{\mathrm{sel}}_{\widehat{ma/m}} \\ & + d\sigma_{m,\mathrm{e}}\left[1 + 2\,(c-1)\,(1-\sigma_{\mathrm{e}})\right] D^{\mathrm{sel}}_{m,\widehat{a/m}} \\ & - c\left[d\sigma_{m,\mathrm{e}} + d\sigma_{m,m,\mathrm{e}}\,(1-2p_m)\right] D^{\mathrm{sel}}_{ma,m} \end{aligned} \tag{B30}$$

$$\begin{aligned} D_{ma}{}' \approx\; & (1-\sigma_{\mathrm{e}} r_{ma})\,D^{\mathrm{sel}}_{ma} + \sigma_{\mathrm{e}} r_{ma}\,D^{\mathrm{sel}}_{m,a} \\ & + 2d\sigma_{m,\mathrm{e}}\,(c-1)\left[(1-\sigma_{\mathrm{e}} r_{ma})\,D^{\mathrm{sel}}_{\widehat{ma/m}} + \sigma_{\mathrm{e}} r_{ma}\,D^{\mathrm{sel}}_{m,\widehat{a/m}}\right] \\ & - (c-1)\left[d\sigma_{m,\mathrm{e}} + d\sigma_{m,m,\mathrm{e}}\,(1-2p_m)\right] D^{\mathrm{sel}}_{ma,m} \end{aligned} \tag{B31}$$

with:

$$D^{\rm sel}_{m\widehat{/}a} \approx (1 - sh)\, D_{m\widehat{/}a} - sh\left(D_{ma\widehat{/}a} + D_{m,a\widehat{/}a}\right) \tag{B32}$$

$$D^{\rm sel}_{m,a} \approx (1 - sh)\, D_{m,a} - s\,(1 - h)\, D_{ma,a} + 2sh\, D_{m,a\widehat{/}a} \tag{B33}$$

$$D^{\rm sel}_{ma} \approx (1 - sh)\, D_{ma} - s\,(1 - h)\, D_{ma,a} + 2sh\, D_{ma\widehat{/}a} \tag{B34}$$

$$D^{\rm sel}_{ma\widehat{/}m} \approx (1 - sh)\, D_{ma\widehat{/}m} - sh\left(D_{ma\widehat{/}ma} + D_{ma\widehat{/}m,a}\right) \tag{B35}$$

$$D^{\rm sel}_{m,a\widehat{/}m} \approx (1 - sh)\, D_{m,a\widehat{/}m} - sh\left(D_{ma\widehat{/}m,a} + D_{m,a\widehat{/}m,a}\right) \tag{B36}$$

$$D^{\rm sel}_{ma,m} \approx (1 - sh)\, D_{ma,m} - s\,(1 - h)\, D_{ma,ma}\,. \tag{B37}$$

When $c = 1$ (no cost of sex) $D_{m\widehat{/}a} = 0$ at quasi-equilibrium, while for $s$, $d$, $\sigma_{\rm e}$ and $r_{ma}$ small and $h_m = 1/2$, $D_{ma} + D_{m,a}$ is approximately:

$$\begin{aligned} D_{ma} + D_{m,a} \approx & -\frac{d\sigma_{m,{\rm e}}\,(2d + 2sh + \sigma_{\rm e})\, T_{ma}\, p_m q_m\, p_a}{4N\,(sh + \sigma_{\rm e})\,(sh + \sigma_{\rm e} r_{ma})\,(2d + 2sh + \sigma_{\rm e} + \sigma_{\rm e} r_{ma})} \\ & + \frac{d\sigma_{m,{\rm e}}\, s\,(1 - h)\,\iota\, p_m q_m\, {p_a}^2}{(sh + \sigma_{\rm e})\,(sh + \sigma_{\rm e} r_{ma})\,(s + \sigma_{\rm e})^2} \end{aligned} \tag{B38}$$

with:

$$\begin{aligned} T_{ma} = & \frac{(sh)^2}{(d + sh + \sigma_{\rm e})\,(2d + sh + \sigma_{\rm e})} + \frac{(sh)^2}{(d + sh + \sigma_{\rm e} r_{ma})\,(2d + sh + \sigma_{\rm e} r_{ma})} \\ & + \frac{4s^2\,(1 - h)\,(d + sh)\,(2sh + \sigma_{\rm e})}{(s + \sigma_{\rm e})^2\,(d + sh + \sigma_{\rm e} r_{ma})\,(2d + 2sh + \sigma_{\rm e} r_{ma})} \\ & - \frac{sh\sigma_{\rm e}\,[2s\,(1 - h)\,d + sh\,[s\,(2 - h) + \sigma_{\rm e}]]}{(s + \sigma_{\rm e})\,(sh + \sigma_{\rm e})\,(d + sh + \sigma_{\rm e} r_{ma})\,(2d + sh + \sigma_{\rm e} r_{ma})}\,. \end{aligned} \tag{B39}$$

Approximations take more complicated forms when $c > 1$; however, expressions for $D_{ma}$, $D_{m,a}$ and $D_{m\widehat{/}a}$ at quasi-equilibrium can be obtained from equations B29 – B37 (see *Mathematica* notebook available as Supplementary Material).

The association $D_{m\widehat{/}a}$ measures the covariance between $p_{m(i)}$ and $p_{a(i)}$ across demes. As shown by equations B29 and B32 (and equations B16–B17, B22–B23 above), it is positive at equilibrium when $c > 1$ (*i.e.*, in the presence of direct selection against allele $m$), and ultimately generated by the variance in linkage disequilibrium across demes. Indeed, the frequency of $m$ and $a$ within a deme both tend to decrease when $D_{ma(i)} > 0$ (due to the fact that both $m$ and $a$ are associated with a deleterious allele at the other locus), while $p_{m(i)}$ and $p_{a(i)}$ tend to increase when $D_{ma(i)} < 0$, causing a positive covariance between $p_{m(i)}$ and $p_{a(i)}$.

Associations $D_{ma}$ and $D_{m,a}$ represent the association between alleles $m$ and $a$ (at the metapopulation scale), either on the same haplotype ($D_{ma}$) or on different haplotypes of the same individual ($D_{m,a}$). When $c > 1$, direct selection occurs against alleles $m$ and $a$, and population structure generates interference between the two loci. However, interference (whose effect is represented by terms in $c - 1$ in equations B30–B31 and in the associations involved) may generate either positive or negative $D_{ma}$ and $D_{m,a}$ depending on parameter values. The variance in linkage disequilibrium among demes tends to generate negative $D_{ma}$ and $D_{m,a}$ through the terms in $D_{ma\widehat{/}m}$, $D_{m,a\widehat{/}m}$, $D_{ma\widehat{/}a}$ and $D_{m,a\widehat{/}a}$ that appear in equations B30–B31 and B33–B34. This corresponds to the classical Hill-Robertson effect (Hill and Robertson, 1966), generating negative LD between deleterious alleles. However, interference also tends to generate negative values of $D_{ma,a}$ and $D_{ma,m}$ (which is both a consequence of the covariance in homozygosity among loci $D_{ma,ma}$ and of the variance in LD among demes), which in turn lead to positive $D_{ma}$ and $D_{m,a}$. Indeed, negative $D_{ma,a}$ and $D_{ma,m}$ mean that alleles $m$ and $a$ tend to be found relatively more often than alleles $M$ and $A$ in heterozygotes at the other locus. Because selection is less efficient among heterozygotes than among homozygotes, selection against allele $a$ tends to be less efficient among individuals carrying allele $m$, leading to a higher frequency of $a$ among those individuals and thus to positive $D_{ma}$, $D_{m,a}$. Furthermore, a second source of positive $D_{ma}$ and $D_{m,a}$ stems from the positive correlation between the frequencies of $m$ and $a$ in uniting gametes within demes (term in $D_{m\widehat{/}a}$ in equation B30). The effect of interference between selected loci in spatially structured populations will be discussed further in the context of our three-locus model.

When $c = 1$ (no direct selection at the modifier locus), these interference effects disappear, but $D_{ma}$ and $D_{m,a}$ are still generated due to the modifier effect. Indeed, the modifier effect generates positive $D_{ma,a}$ (when $m$ increases investment in sex), so that allele $m$ tends to be found relatively more often in homozygotes at the selected locus, among which selection against allele $a$ is more efficient. This generates a negative association between $m$ and $a$ (negative $D_{ma}$ and $D_{m,a}$). The associations $D_{ma\widehat{/}m}$ and $D_{m,a\widehat{/}m}$ (which are negative) also tend to generate negative $D_{m,a}$ even when $c = 1$ (as shown by equation B30). Indeed, when $m$ is locally associated with $a$ ($D_{ma(i)} > 0$, $D_{m,a(i)} > 0$), the frequency of $m$ tends to decrease locally. Therefore, gametes carrying

$m$ have a relatively higher chance of fusing with a gamete carrying $M$, and thus $A$ (since $D_{ma(i)} > 0$), generating a negative association between $m$ and $a$ on different haplotypes (negative $D_{m,a}$). Conversely, when $m$ is locally associated with $A$, the frequency of $m$ tends to increase locally, and gametes carrying $m$ have a relatively higher chance of fusing with another gamete carrying $m$, and thus $A$ (also generating negative $D_{m,a}$). This stochastic mechanism that tends to favor sex (by generating negative associations between alleles increasing the rate of sex and deleterious alleles at other loci) has been described by Roze (2014) in the case of a haploid model. Finally, the association $D_{ma,m}$ (which is negative) has the opposite effect and tends to generate positive $D_{m,a}$ even when $c = 1$. Negative $D_{ma,m}$ means that allele $A$ is relatively more frequent among homozygotes at the modifier locus than among heterozygotes. Because $mm$ homozygotes have sex more frequently with $Mm$ heterozygotes (in which the frequency of $a$ is relatively higher) than $MM$ homozygotes, this generates a positive associations between $m$ and $a$ on different haplotypes (positive $D_{m,a}$). Overall, one can show that when $c = 1$ (no cost of sex), the effect of $D_{ma,m}$ (generating positive $D_{m,a}$) is weaker than the combined effects of $D_{ma\widehat{/}m}$, $D_{m,a\widehat{/}m}$ and $D_{ma,a}$ (generating negative $D_{m,a}$), so that $D_{ma}$ and $D_{m,a}$ are negative at equilibrium. When $c > 1$, however, $D_{ma}$ and $D_{m,a}$ may become positive, in particular when the rate of sex is large.
