## Supplementary material for "Deleterious mutations and selection for sex in spatially structured, diploid populations": File S3

### FILE S3: THREE-LOCUS MODEL

Our model was extended to include a second selected locus with two alleles $B$ and $b$, where $b$ is deleterious (with the same selection and dominance coefficients $s$ and $h$ as allele $a$ at the first selected locus), assuming that selection is multiplicative across loci (no epistasis). The recombination rate between the modifier and this second selected locus is denoted $r_{mb}$, while the recombination rate between the two selected loci is denoted $r_{ab}$. We provide here recursions for genetic associations between the two selected loci, while expressions for three-locus associations are derived in the *Mathematica* notebook available as Supplementary Material. Expressing indirect selection for sex generated by the interaction between the two selected loci required computing a large number of such three-locus associations, which often take complicated forms. This three-locus model can then be extrapolated to the case of a linear chromosome (along which deleterious mutations occur at a rate $U$ per generation, at an infinite number of possible sites). As explained below, this involves integrating numerically the result from the three-locus model over all possible positions of deleterious alleles along the chromosome. As shown in the Supplementary Material, simpler expressions can be obtained by assuming that all loci recombine freely and that $s$, $d$ and $\sigma_{\mathrm{e}}$ are small. For the parameter values used in the paper, and when the average number of crossovers per chromosome is sufficiently large, we found that the evolutionarily stable rate of sex predicted from these simpler expressions is often very close to the result obtained by integrating numerically the expressions obtained under arbitrary recombinations rates, $d$ and $\sigma_{\mathrm{e}}$.

**Associations between the two selected loci.** Associations $D_{ab,ab}$, $D_{ab\widehat{/}ab}$, $D_{ab\widehat{/}a,b}$ and $D_{a,b\widehat{/}a,b}$ are generated by population structure. Recursions take similar forms as for the equivalent associations between the modifier and the first selected locus (see File S2, which also provides interpretations for these associations):

$$D_{a,b\widehat{/}a,b}{}' \approx \frac{p_a\, p_b}{2N} + (1-d)^2\,(1-\sigma_{\mathrm{e}})^2\,(1-4sh)\, D_{a,b\widehat{/}a,b} \tag{C1}$$

$$D_{ab\widehat{/}a,b}{}' \approx (1-d)^2\,(1-\sigma_{\mathrm{e}})\,(1-4sh)\left[(1-\sigma_{\mathrm{e}} r_{ab})\, D_{ab\widehat{/}a,b} + \sigma_{\mathrm{e}} r_{ab}\, D_{a,b\widehat{/}a,b}\right] \tag{C2}$$

$$D_{ab\overset{\frown}{/}ab}{}' \approx \frac{p_a\,p_b}{2N} + (1-d)^2\,(1-4sh)\,\Big[(1-\sigma_{\mathrm{e}}\,r_{ab})^2\,D_{ab\overset{\frown}{/}ab} + 2\sigma_{\mathrm{e}}\,r_{ab}\,(1-\sigma_{\mathrm{e}}\,r_{ab})^2\,D_{ab\overset{\frown}{/}a,b} + (\sigma_{\mathrm{e}}\,r_{ab})^2\,D_{a,b\overset{\frown}{/}a,b}\Big] \tag{C3}$$

$$D_{ab,ab}{}' \approx (1-\sigma_{\mathrm{e}})\,(1-2s)\,D_{ab,ab} + \sigma_{\mathrm{e}}\,(1-4sh)\,\Big[(1-r_{ab})^2\,D_{ab\overset{\frown}{/}ab} + 2r_{ab}\,(1-r_{ab})^2\,D_{ab\overset{\frown}{/}a,b} + r_{ab}{}^2 D_{a,b\overset{\frown}{/}a,b}\Big]. \tag{C4}$$

When $s$, $d$ and $\sigma_{\mathrm{e}}$ are small, approximations for these associations at equilibrium are given by equations B12 – B15 in File S2, replacing $s$ by $2s$, $r_{ma}$ by $r_{ab}$ and $p_m q_m$ by $p_b$.

Associations $D_{ab\overset{\frown}{/}a}$, $D_{a,b\overset{\frown}{/}a}$ and $D_{ab,a}$ are generated by the previous associations and by the effect of selection. Again, recursions take similar forms as the recursions for $D_{ma\overset{\frown}{/}m}$, $D_{m,a\overset{\frown}{/}m}$ and $D_{ma,m}$ in File S2:

$$D_{a,b\overset{\frown}{/}a}{}' \approx (1-d)^2\,(1-\sigma_{\mathrm{e}})\,\Big[(1-3sh)\,D_{a,b\overset{\frown}{/}a} - sh\,\Big(D_{ab\overset{\frown}{/}a,b} + D_{a,b\overset{\frown}{/}a,b}\Big)\Big] \tag{C5}$$

$$D_{ab\overset{\frown}{/}a}{}' \approx (1-d)^2\,\Big[(1-3sh)\,\Big[(1-\sigma_{\mathrm{e}}\,r_{ab})\,D_{ab\overset{\frown}{/}a} + \sigma_{\mathrm{e}}\,r_{ab}\,D_{a,b\overset{\frown}{/}a}\Big] - sh\,\Big[(1-\sigma_{\mathrm{e}}\,r_{ab})\,D_{ab\overset{\frown}{/}ab} + D_{ab\overset{\frown}{/}a,b} + \sigma_{\mathrm{e}}\,r_{ab}\,D_{a,b\overset{\frown}{/}a,b}\Big]\Big] \tag{C6}$$

$$D_{ab,a}{}' \approx (1-\sigma_{\mathrm{e}})\,[(1-s-sh)\,D_{ab,a} - s\,(1-h)\,D_{ab,ab}] + \sigma_{\mathrm{e}}\,(1-3sh)\,\Big[(1-r_{ab})\,D_{ab\overset{\frown}{/}a} + r_{ab}\,D_{a,b\overset{\frown}{/}a}\Big] - \sigma_{\mathrm{e}}\,sh\,\Big[(1-r_{ab})\,D_{ab\overset{\frown}{/}ab} + D_{ab\overset{\frown}{/}a,b} + r_{ab}\,D_{a,b\overset{\frown}{/}a,b}\Big]. \tag{C7}$$

As explained in File S2, $D_{ab\overset{\frown}{/}a}$ and $D_{a,b\overset{\frown}{/}a}$ are negative at equilibrium, reflecting a negative covariance between the frequency of $a$ within demes and the genetic association between $a$ and $b$ (either on the same or on different chromosomes of an individual) within the deme: indeed, $a$ tends to decrease in frequency when it is associated with $b$, since $b$ is deleterious. $D_{ab,a}$ is also negative at equilibrium, reflecting the fact that the frequency of $b$ tends to be higher in heterozygotes at the first selected locus than in homozygotes. This is due to the fact that selection against $b$ is more efficient among homozygotes at the first selected locus than among heterozygotes (due to the correlation in homozygosity among loci represented by $D_{ab,ab}$), and also to the negative covariance between the linkage disequilibrium between $a$ and $b$ and the frequency of $a$ among gametes (see File S2). Associations $D_{ab\overset{\frown}{/}b}$, $D_{a,b\overset{\frown}{/}b}$ and $D_{ab,b}$ are generated by the same mechanisms and take the same forms.

Finally, recursions on pairwise associations $D_{a\overset{\frown}{/}b}$, $D_{a,b}$ and $D_{ab}$ are given by:

$$D_{a\overset{\frown}{/}b}{}' \approx (1-d)^2\, D^{\mathrm{sel}}_{a\overset{\frown}{/}b} \tag{C8}$$

$$D_{a,b}{}' \approx (1-\sigma_{\mathrm{e}})\, D^{\mathrm{sel}}_{a,b} + \sigma_{\mathrm{e}}\, D^{\mathrm{sel}}_{a\overset{\frown}{/}b} \tag{C9}$$

$$D_{ab}{}' \approx (1-\sigma_{\mathrm{e}} r_{ab})\, D^{\mathrm{sel}}_{ab} + \sigma_{\mathrm{e}} r_{ab}\, D^{\mathrm{sel}}_{a,b} \tag{C10}$$

where associations after selection are given by:

$$D^{\mathrm{sel}}_{a\overset{\frown}{/}b} \approx (1-2sh)\, D_{a\overset{\frown}{/}b} - sh\left(D_{ab\overset{\frown}{/}a} + D_{a,b\overset{\frown}{/}a} + D_{ab\overset{\frown}{/}b} + D_{a,b\overset{\frown}{/}b}\right) \tag{C11}$$

$$D^{\mathrm{sel}}_{a,b} \approx (1-2sh)\, D_{a,b} + 2sh\left(D_{a,b\overset{\frown}{/}a} + D_{a,b\overset{\frown}{/}b}\right) - s\,(1-h)\,(D_{ab,a} + D_{ab,b}) \tag{C12}$$

$$D^{\mathrm{sel}}_{ab} \approx (1-2sh)\, D_{ab} + 2sh\left(D_{ab\overset{\frown}{/}a} + D_{ab\overset{\frown}{/}b}\right) - s\,(1-h)\,(D_{ab,a} + D_{ab,b})\,. \tag{C13}$$

Equations C8 and C11 show that population structure and selection tend to generate a positive covariance between the frequencies of $a$ and $b$ within demes (measured by the association $D_{a\overset{\frown}{/}b}$). This is due to the variance in linkage disequilibrium (and in the association across haplotypes $D_{a,b}$) between demes: selection against deleterious alleles is more efficient in demes where LD is positive (since $a$ and $b$ tend to be present in the same individuals), reducing the frequency of $a$ and $b$ in those demes, while selection is less efficient in demes where LD is negative, increasing the frequency of $a$ and $b$ in those demes. The positive covariance between the frequencies of $a$ and $b$ in gametes sampled from the same deme generates a positive association between $a$ and $b$ on the two chromosomes of the same individual, $D_{a,b}$ (second term of equation C9), which contributes to generating positive $D_{ab}$ (as $D_{a,b}$ is converted into $D_{ab}$ by recombination — second term of equation C10). As shown by equations C12 – C13, a second source of positive $D_{a,b}$ and $D_{ab}$ involves the associations $D_{ab,a}$, $D_{ab,b}$ (which are negative, as explained above). Again, a negative $D_{ab,a}$ means that the frequency of $b$ is higher among $Aa$ heterozygotes than among $AA$, $aa$ homozygotes. Since selection against $a$ is more efficient among homozygotes than among heterozygotes, this generates a positive association between $a$ and $b$ (either on the same or on different chromosomes of the same individual). These two different sources of positive $D_{ab}$ and $D_{a,b}$ (through $D_{a\overset{\frown}{/}b}$, and through $D_{ab,a}$, $D_{ab,b}$) are opposed by a mechanism that contributes to generate negative $D_{ab}$ and $D_{a,b}$, represented by the terms in $2sh$ in equations C12 and C13. This corresponds to the classical Hill-Robertson effect (Hill and Robertson, 1966; Felsenstein, 1974; Martin et al., 2006): selection against $a$ and $b$ is more efficient within demes where LD (and the association across haplotypes) is positive, which decreases the frequencies of the deleterious alleles and the magnitude of LD in those demes, while selection is less efficient in demes where LD is negative, increasing the frequencies of $a$ and $b$ and the magnitude of LD in those demes. These effects on allele frequencies within demes thus tend to generate negative $D_{ab}$ and $D_{a,b}$ at the scale of the metapopulation.

Under obligate sex and complete dispersal ($\sigma_{\mathrm{e}} = 1$, $d = 1$), $D_{ab}$ is positive and given by:

$$D_{ab} \approx \frac{s^2 h \left(1-h\right) p_a p_b}{N \left(r_{ab} + 2sh\right)}. \tag{C14}$$

Still under obligate sex ($\sigma_{\mathrm{e}} = 1$) and when $d$, $s$ and $r_{ab}$ are small, one obtains:

$$D_{ab} \approx \frac{s^2 h \left[h r_{ab} + 2\left(1-3h\right)\left(d+sh\right)\right] p_a p_b}{4N \left(d+sh\right)\left(2sh + r_{ab}\right)\left(d + 2sh + r_{ab}\right)\left(2d + 3sh + r_{ab}\right)}. \tag{C15}$$

From this, one predicts that $D_{ab}$ should be always positive when $h < 1/3$, while it may become negative when $h > 1/3$. Finally, when $\sigma_{\mathrm{e}}$, $d$, $s$ and $r_{ab}$ are small, we have:

$$D_{ab} \approx \frac{s^2 \left[h\left(1-3h\right){\sigma_{\mathrm{e}}}^2 + \left[2\left(1-h\right)^2 d + \left[5 - h\left(14-h\right)\right] sh\right]\sigma_{\mathrm{e}} - 4\left(1+h\right)s^2h^2\right] p_a p_b}{2N\left(2s+\sigma_{\mathrm{e}}\right)\left(s+sh+\sigma_{\mathrm{e}}\right)\left(2sh + r_{ab}\sigma_{\mathrm{e}}\right)\left(d+2sh+r_{ab}\sigma_{\mathrm{e}}\right)\left(2d+3sh+r_{ab}\sigma_{\mathrm{e}}\right)}. \tag{C16}$$

The association $D_{ab}$ measures the linkage disequilibrium between the deleterious alleles at the whole metapopulation level. One can show that the average LD between $a$ and $b$ within demes, defined as:

$$\mathrm{E}_i\left[D_{ab(i)}\right] = \frac{1}{2}\mathrm{E}_{ij}\left[\left(p_{a(ij1)} - p_{a(i)}\right)\left(p_{b(ij1)} - p_{b(i)}\right) + \left(p_{a(ij2)} - p_{a(i)}\right)\left(p_{b(ij2)} - p_{b(i)}\right)\right] \tag{C17}$$

is equivalent to $D_{ab} - D_{\widehat{a/b}}$. Under obligate sex ($\sigma_{\mathrm{e}} = 1$) and when $d$, $s$ and $r_{ab}$ are small, this is approximately:

$$D_{ab} - D_{\widehat{a/b}} \approx \frac{s^2 h \left[\left(1-3h\right)d + \left(1-4h\right)sh\right] p_a p_b}{2N\left(d+sh\right)\left(2sh+r_{ab}\right)\left(d+2sh+r_{ab}\right)\left(2d+3sh+r_{ab}\right)}. \tag{C18}$$

When the dispersal rate $d$ tends to zero, equation C18 converges to the result obtained by Roze (2021) in the case of a single finite population. Furthermore, when the terms in $D_{ab,a}$, $D_{ab,b}$ are removed from equations C12 – C13, $D_{ab} - D_{\widehat{a/b}}$ becomes always negative, indicating that positive LD within demes is only due to those terms.

**Three-locus model, and extrapolation to the case of a linear chromosome.** The change in allele frequencies at the sex modifier locus in the three-locus model is generated by a large number of genetic associations between the three loci. Some of these associations are generated by population structure (e.g., $D_{mab\overset{\frown}{/}mab}$, $D_{mab,mab}$, ...), others by population structure and selection against deleterious alleles (e.g., $D_{mab\overset{\frown}{/}ma}$, $D_{mab,ma}$, ...), others by population structure and the effect of the sex modifier (e.g., $D_{mab\overset{\frown}{/}ab}$, $D_{mab,ab}$, ...), and yet others by population structure, selection against deleterious alleles and the sex modifier (e.g., $D_{mab\overset{\frown}{/}a}$, $D_{mab}$, ...). Furthermore, associations between the modifier and the first selected locus (derived in File S2) are affected by the second selected locus. Although it should be possible in principle to provide an explanation for all the different effects generating these associations, we do not attempt to do that here due to the very large number of associations involved. Recursions for associations can be computed automatically using the *Mathematica* notebook available as Supplementary Material, for arbitrary recombination rates among the three loci. This yields complicated expressions for associations at quasi-linkage equilibrium, which can then be integrated numerically over all possible positions of selected loci along the chromosome, in order to obtain the net strength of selection at the sex modifier locus when deleterious mutations may occur at a large number of possible sites, assuming a chromosome map length $R$ and a uniform density of crossovers along the chromosome. This numerical integration is computationally demanding, however, and we observed that when the chromosome map length is sufficiently large ($R = 10$ in our simulations, in order to mimic a genome with multiple chromosomes), the predicted evolutionarily stable rate of sex is often close to the result obtained using simpler recursions assuming free recombination among all loci and weak rates of sex and dispersal (these recursions are also given in the Supplementary Material). To leading order, one obtains that the change in frequency of allele $m$ is given by:

$$\Delta p_m \approx s_{\text{direct}} + s_{\text{indirect}} \tag{C19}$$

with (assuming $h_m = 1/2$ for simplicity):

$$s_{\text{direct}} \approx -(c-1)\, d\sigma_{m,\text{e}} \left(p_m q_m + D_{m,m} - 2D_{m\overset{\frown}{/}m}\right), \tag{C20}$$

$$s_{\text{indirect}} \approx -sh \sum_a \left(D_{ma} + D_{m,a} - 2D_{m\overset{\frown}{/}a}\right) - s\,(1-2h) \sum_a D_{ma,a} \tag{C21}$$

where the sums in equation C21 are over all selected loci. Under free recombination among all loci, equation C21 takes the form:

$$s_{\text{indirect}} \approx \frac{1}{N}\left[F_1\left(s,\, h,\, d,\, \sigma_{\text{e}},\, c\right)\sum_a p_a + F_2\left(s,\, h,\, d,\, \sigma_{\text{e}},\, c\right)\sum_a p_a \sum_b p_b\right] p_m q_m \quad \text{(C22)}$$

where $F_1$ and $F_2$ are complicated functions of the parameters obtained from the two- and three-locus models (respectively), while $\sum_a p_a \approx U/\left(sh\right)$ under our assumptions $(Ns,\ Nd \gg 1)$. The first term between brackets in equation C22 is thus of order $U$, while the second is of order $U^2$. Because the associations $D_{m,m}$ and $D_{m\widehat{/}m}$ are affected by selected loci, their expressions at QLE also involve terms in $U$ and $U^2$. For given values of $N$, $d$, $s$, $h$, $U$ and $c$, the evolutionarily stable rate of sex can be obtained by computing $\Delta p_m$ for a range of values of $\sigma_{\text{e}}$, and find the value of $\sigma_{\text{e}}$ for which $\Delta p_m = 0$ by interpolation (see Supplementary Material).
