## Supplementary material for "Deleterious mutations and selection for sex in spatially structured, diploid populations": Figure S1

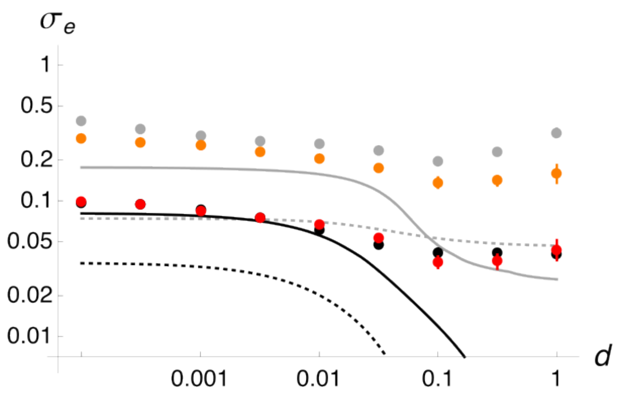


**Figure S1.** Same as Figure 6C, with additional simulation results with a mutation rate $\mu = 10^{-5}$ at the modifier locus, for $c = 1$ (orange dots) and $c = 1.2$ (red dots). Grey and black dots are the same as in Figure 6C and correspond to simulation results obtained with $\mu = 10^{-4}$.
